# Longitudinal bioluminescent imaging of HIV-1 infection during antiretroviral therapy and treatment interruption in humanized mice

**DOI:** 10.1101/745125

**Authors:** John D. Ventura, Jagadish Beloor, Edward Allen, Tongyu Zhang, Kelsey A. Haugh, Pradeep D. Uchil, Christina Ochsenbauer, Collin Kieffer, Priti Kumar, Thomas J. Hope, Walther Mothes

**Affiliations:** Department of Microbial Pathogenesis, Yale University School of Medicine, New Haven, CT 06536, USA; Department of Internal Medicine, Yale University School of Medicine, New Haven, CT 06510, USA; Department of Cellular and Molecular Biology, Feinberg School of Medicine, Northwestern University, Chicago, IL 60611, USA; School of Molecular and Cellular Biology, College of Liberal Arts and Sciences, University of Illinois at Urbana-Campaign, Urbana, IL 61801, USA; Department of Medicine, University of Alabama at Birmingham, Birmingham, Alabama, AL 35233, USA

## Abstract

Non-invasive bioluminescent imaging (NIBLI) of HIV-1 infection dynamics allows for real-time monitoring of viral spread and the localization of infected cell populations in living animals. In this report, we describe full-length replication-competent GFP and Nanoluciferase (Nluc) expressing HIV-1 reporter viruses from two clinical transmitted / founder (T/F) stains: TRJO.c and Q23.BG505. By infecting humanized mice with these HIV-1 T/F reporter viruses, we were able to directly monitor longitudinal viral spread at whole-animal resolution via NIBLI at a sensitivity of as few as 30-50 infected cells. Bioluminescent signal strongly correlated with HIV-1 infection and responded proportionally to virus suppression in vivo in animals treated daily with a combination antiretroviral therapy (cART) regimen. Longitudinal NIBLI following cART withdrawal visualized tissue-sites that harbored virus during infection recrudescence. Notably, we observed rebounding infection in the same lymphoid tissues where infection was first observed prior to ART treatment. Our work demonstrates the utility of our system for studying in vivo viral infection dynamics and identifying infected tissue regions for subsequent analyses.

**Author Summary:** Non-invasive bioluminescent imaging (NIBLI) in small animals allows for in vivo longitudinal imaging of infection spread and pathogenesis. We have taken advantage of the small luciferase reporter protein, Nanoluciferase (Nluc), to generate a replication-competent HIV-1 reporter virus to allow for NIBLI of viral infection in humanize mice. NIBLI via Nluc enabled us to directly visualize longitudinal spreading patterns before, during, and after interruption of daily doses of combined antiretroviral therapy (cART). We observed that rebounding infection often emerged in tissue regions originally associated with infected cells prior to cART treatment. Thus, Nluc-based NIBLI of HIV-1 infection can be used as an experimental tool to study early events involved in viral dissemination and spread from initial sites of infection to draining lymphoid tissues as well as locate infected tissues for subsequent cellular characterization of HIV-1 infected cells.

## Introduction

Whole-body non-invasive bioluminescent imaging (NIBLI) of bacterial and viral pathogens in small animal models is a powerful and versatile experimental tool. NIBLI enables longitudinal real-time monitoring of infection in same animal by exploiting the high sensitivity and dynamic range offered by bioluminescent reporter genes such as luciferases. As a result, critical factors related to pathogenicity, transmissibility, and in vivo dissemination can be elucidated that may go unnoticed in in vitro experimental systems (1, 2). Accordingly, bioluminescence reporter constructs have been developed for longitudinal NIBLI of several pathogens, including influenza, the plaque bacterium *Yersinia pestis*, *Escherichia coli*, methicillin-resistant *Staphylococcus aureus* (MRSA), vaccinia virus, herpes simplex virus-1 (HSV-1), the smallpox virus *Variola major*, and the alphaviruses Sindbis, Chikungunya, Eastern and Venezuelan equine encephalitis viruses (3–12).

Currently, there is great interest in extending non-invasive imaging technology to longitudinally image HIV-1 and SIV infection in vivo. Position Emission Tomography (PET) using Copper (^64^Cu)-labeled α-SIV Gp120 immune-labeling of SIV_mac239_ (i.e. immuno-PET) allows non-invasive real-time monitoring of infected non-human primates (NHPs) by detecting virus compartmentalization in living animals (13). Given the high costs of immuno-PET imaging in NHPs, non-PET based NIBLI systems for small animal models would be more widely accessible and complement the immune-PET imaging in NHP. Therefore, the development of NIBLI for HIV-1 in humanized mice would be an asset for investigators interested in viral transmission, pathogenesis, and whole-animal spread.

NIBLI of longitudinal HIV-1 infection would require the specific and stable labeling of productively infected cells with a bioluminescent reporter detectable in vivo. The small 19 kDa luciferase Nanoluciferase (Nluc) emerges as an ideal candidate due to its very short gene length and its 150-fold brighter luminal capacity than both firefly and *Renilla* luciferases (1, 14). Furthermore, replication-competent Nluc based reporters have already been successfully used in in vivo bioluminescent imaging of longitudinal influenza infection, testifying to the enhanced in vivo functionality, stability, and overall tractability of the Nluc reporter gene (11, 12).

In this report, we developed and characterized full-length replication-competent Nanoluciferase (Nluc) and GFP expressing HIV-1 reporter viruses from two clinical T/F stains, TRJO.c and Q23.BG505. Our design introduces Nluc or GFP at the native *nef* locus upstream of either a full-length or attenuated encephalomyocarditis virus (ECMV) internal ribosome entry site (IRES) sequence to maintain Nef expression (Fig. 1A). The simple insertion of an IRES element produces a bicistronic *gfp* and *nef* transcript that initiates at the native *nef* start codon without additional modification to the genome. Consequentially, the IRES element drives Nef protein expression to near wild-type levels and preserves its function. Also, virus particle infectivity was minimally compromised by the introduction of the reporter gene upstream of the attenuated ECMV IRES element 6ATRi. Reporter gene expression was stable for over 10 days during progressive infection in cultured primary CD4^+^ T cells and for at least two weeks in humanized PBMC mice (Hu-PBL). Nluc expressing T/F reporter viruses permitted NIBLI of longitudinal HIV-1 dissemination from initial infection sites to subsequent systemic spread in multiple humanized mouse models with a sensitivity of as few as 30-50 infected cells. Nluc signal was strongly correlated with both the increase of reverse transcriptase (RT) activity and the number of infected cells in blood plasma. Finally, we demonstrate the utility of our system for localizing and subsequently studying HIV-1 tissue reservoirs involved in establishing recrudescent infection following cART withdrawal.

**Figure 1.**
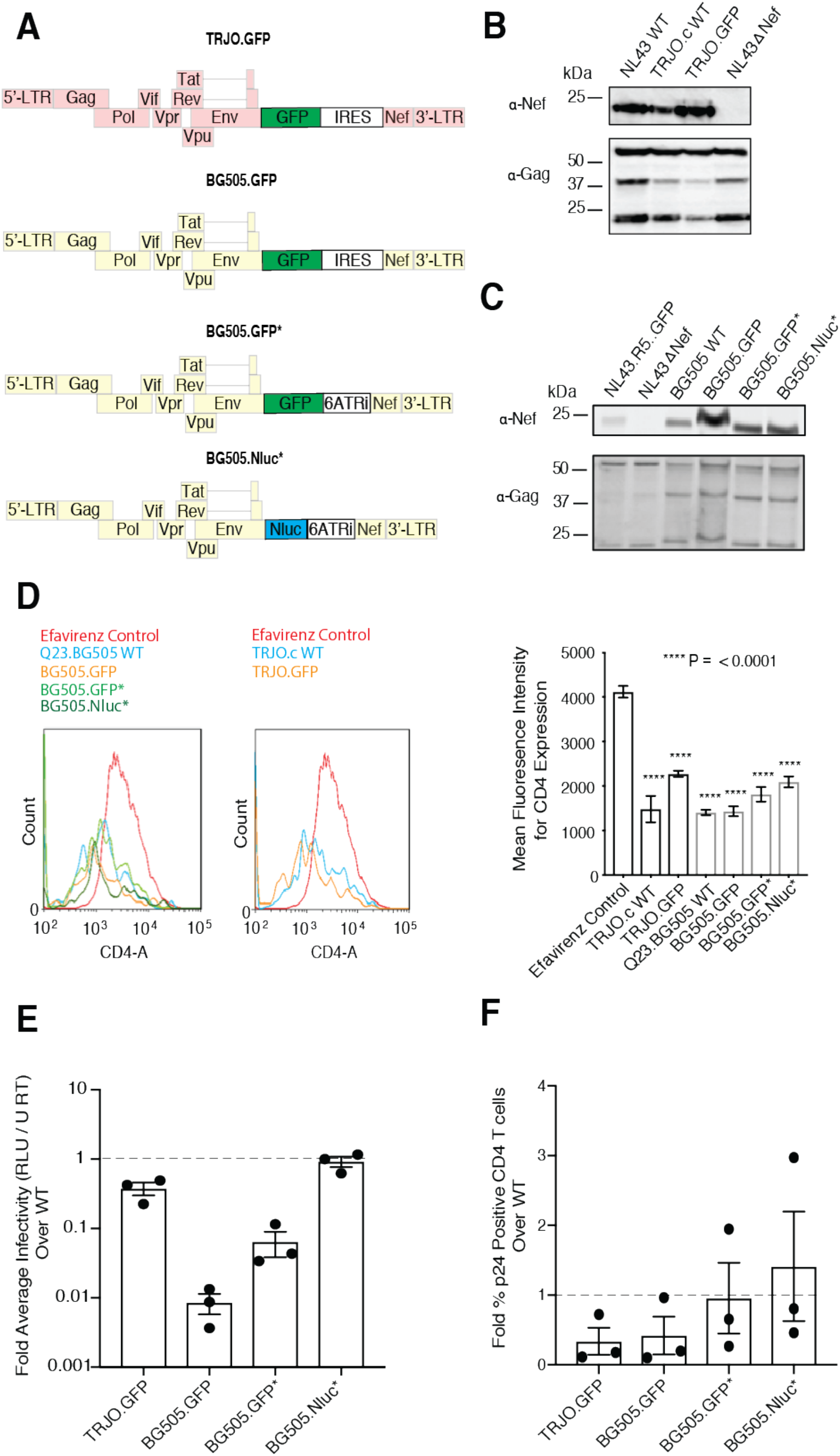
Functional characterization of HIV-1 T/F full-length reporter viruses. (A) HIV-1 T/F full-length reporter virus design. (B-C) Western blot analysis of Nef protein expression in HEK293 producer cell lysates transfected with TRJO.c (B) and Q23.BG505 (C) derived T/F full-length reporter virus plasmid DNA. Equal amounts of p55 Gag were loaded onto each lane to assess the relative Nef expression. (D) Surface CD4 surface expression on p24+ Jurkat CCR5+ JLTRGFP.R5 cells infected with HIV-1 T/F wild-type or HIV-1 T/F reporter virus 48 hours after initiating infection. Data shown as mean +/-SD of 3 technical replicates with significance calculated from a one-way ANOVA. (E) Single-round infectivity of HIV-1 T/F reporter viruses on TZM-bl cells cells in the presence of 10 nM of the protease inhibitor saquinavir to block virus spreading (n=3). Infectivity for each virus was determined by measuring the fold firefly luciferase expression from infected TZM-bl cells above uninfected negative controls and normalizing to reverse transcriptase units (RT U). Data displayed is average fold HIV-1 T/F reporter virus infectivity over TRJO.c and Q23.BG505 wild-type virus +/-SEM from three independent experiments. (F) HIV-1 T/F reporter virus spread in primary CD4+ T cells (n=3). Each sample was set to the same level of p24 positive cells and then allowed to spread for four days in culture. Data displayed as the fold HIV-1 T/F reporter virus infection (as the value of % p24) over wild-type +/-the SEM from three independent CD4+ T cell preparations.

## Results

### Reporter virus design to label productively infected cells

We designed GFP and Nluc HIV-1 reporter viruses from two clinical T/F HIV-1 strains: TRJO.c, a sexually-transmitted clade B strain (15), and the Q23.BG505 clade A strain carrying the BG505 envelope protein of a vertically transmitted virus in the clade A backbone of Q23 (16). Reporter gene transcription was designed to initiate at the native start codon for the *nef* open reading frame and expresses the reporter gene from an integrated proviral sequence within the genome of a productively infected target cell (Fig 1A). Nef protein expression was driven by introducing one of two different Internal Ribosomal Entry Site (IRES) elements: the wild-type ECMV IRES or a truncated ECMV IRES derivative, 6ATRi that exhibits attenuated levels of gene expression (17). The use of IRES sequences was chosen to avoid defects in Nef functionality shown to occur when self-cleavage peptides such as T2A were placed upstream of the Nef coding region that resulted in the generation of non-myristoylated Nef (17). In total, we constructed four HIV-1 T/F reporter viruses designed to express either GFP or Nluc in productively infected cells: TRJO.GFP.IRES.Nef (hereafter called TRJO.GFP), Q23.BG505.GFP.IRES.Nef, (hereafter called BG505.GFP), Q23.BG505.GFP.6ATRi.Nef (hereafter called BG505.GFP*) and Q23.BG505.Nluc.6ATRi.Nef (hereafter called BG505.Nluc*) (Fig 1A).

### HIV-1 T/F reporter viruses express functional Nef

We first tested whether Nef protein expression and function were affected by the insertion of the both reporter genes and IRES elements. To determine Nef protein expression levels, we transfected HEK293 cells with HIV-1 T/F plasmid DNA and subjected cell lysate to western blot analysis. To assess the relative Nef expression, all samples were normalized to equal amounts of capsid p55 protein. Wild-type TRJO.c virus exhibited slightly weaker Nef expression than the laboratory adapted strain NL4-3. TRJO.GFP containing the wild-type ECMV IRES sequence, restored Nef expression to levels comparable to that of wild-type NL4-3 (Fig. 1B). For Q23.BG505 T/F reporter viruses, we observed a higher expression of Nef from producer HEK293 cell for viruses containing the ECMV IRES sequence, and the attenuated 6ATRi restored Nef expression to levels similar to that of wild-type Q23.BG505 (Fig. 1C).

Nef is a crucial HIV-1 accessory protein required to elicit a pathogenic infection in vivo by, in part, facilitating in the removal of cell surface proteins such as CD4 and Major Histocompatibility Complex Class I from the plasma membrane of infected cells (18–22). We therefore tested Nef function by infecting the Jurkat reporter cell line JLTRGFP.R5 that stably expresses CCR5 and expresses GFP in the presence of HIV-1 Tat (23). This reporter cell line allowed us to isolate cells infected with both wild-type and cognate T/F reporter viruses and compare surface CD4 expression to that of cells treated with the non-nucleoside reverse transcriptase inhibitor Efavirenz. JLTRGFP.R5 cells were infected with the same 50 μl volume of HIV-1 wild-type or HIV-1 T/F reporter viruses. All samples were analyzed by p24 intracellular flow cytometry after 48 hours and fresh JLTRGFP.R5 cells were added to ensure each sample had the same ratio of infected: uninfected cells (∼0.6% in this experiment). After 48 additional hours the total CD4 surface expression in the GFP^+^ (i.e. HIV-1 infected) cell population was assessed. CD4 expression was significantly reduced in all reporter virus-infected cell populations when compared to an RT inhibitor control, which included 1 μM efavirenz, and this reduction was comparable to wild-type infected populations (Fig. 1D). Together, these data revealed that each HIV-1 T/F reporter virus maintained Nef expression and functionality.

### Viral infectivity of the 6ATRi IRES containing reporter viruses is not compromised

We next tested whether the introduction of the reporter gene and IRES sequences impaired single-round infectivity. Infectivity was assessed by infecting TZM-bl indicator cells in the presence of 16 μg/ml DEAE-Dextran to ensure efficient virus binding and 10 nM of Saquinavir, a HIV-1 protease inhibitor, to prevent multiple rounds of infection. Infectivity was measured 48 hours later as the fold bioluminescent readout from firefly luciferase from infected cells above control cells treated with 1 μM Efavirenz. These values were normalized to reverse transcriptase activity (RT Units) measured for each viral stock. Single-round TRJO.GFP virus infectivity was reduced by about two-fold compared to TRJO.c wild-type strain infectivity levels (Fig. 1E). Single-round infectivity was more variable for Q23.BG505-derived T/F reporter viruses. The BG505.GFP reporter virus using the native IRES sequence to express Nef exhibited greatly impaired infectivity (∼ 2 logs), possibly as a result of cytotoxic effects caused by the observed Nef overexpression (Fig. 1E). Single-round infectivity for each reporter virus possessing the 6ATRi IRES sequence (i.e. BG505.GFP* and BG505.Nluc*) more closely matched levels observed for the Q23.BG505 parental strain (Fig. 1E).

We next tested the spreading efficiency over a 72-hour period in primary CD4^+^ T cells. Primary human CD4^+^ T cells were isolated from human PBMCs and infected with each HIV-1 T/F reporter virus in parallel with their parental strains. After 48 hours, all infected cell samples were measured for total p24^+^ expression by flow cytometry and all cells were set to the lowest p24^+^ percentage so that each cell preparation contained the same ratio of infected to uninfected cells as a starting point. Virus was allowed to spread for an additional 72 hours and then measured for total HIV-1 infected cells by flow cytometry (Fig. 1F). We observed a two-fold decrease in p24^+^ cells in TRJO.GFP infected samples in comparison to samples infected with the TRJO.c parental strain (Fig. 1F). The T/F reporter viruses BG505.GFP* and BG505.Nluc* again closely matched the behavior of the parental Q23.BG505 virus as observed for the single-round infectivity data making them the best-behaved T/F reporter viruses.

### Relationship between reporter expression and the p24^+^ signal

Considering that our T/F reporter viruses were designed to regulate transcriptional expression of Nluc or GFP via the *nef* start codon, and therefore would be expressed in a similar manner as the *nef* locus (i.e. before the production of HIV-1 capsid) we wanted to experimentally evaluate the relationship between GFP reporter expression and the p24 signal using flow cytometry. CCR5 coreceptor expressing CEM.SS.R5.NKr T lymphocyte cells were infected with 1 x 10^6^ infectious units (IUs) of BG505.GFP and BG505.GFP* T/F reporter virus, and the levels of GFP and p24 expression were measured by p24 intracellular flow cytometry at 4, 8, 18, and 24 hours post-infection. Interestingly, a large fraction of cells immediately stained positive for p24 before productive infection could be established (Supplementary Fig. 1A-C). Cells became p24^+^ even in the presence of the RT inhibitor Efavirenz that blocks infection. This early p24^+^ signal increased until 8 hours post-infection and then declined (Supplementary Fig. 1B-C). This indicates that early virus binding and internalization of HIV particles results in a p24^+^/GFP^-^ signal at the flow cytometry level that is independent of infection. GFP reporter expression was observed beginning at 18 hours post-infection and continued to increase. Consistent with productive infection at that time point, the p24^+^ population again increased in parallel resulting in a robust double-positive GFP^+^/p24^+^ population. Lastly, we observed that the fraction of double-positive cells within the total HIV-1 infected cell population (i.e. total p24^+^ cells) was lower in cells infected with the BG505.GFP virus, which contains the full-length ECMV IRES suggesting that overexpression of Nef may lead to an increase in cellular cytotoxicity, in agreement with our single-round infectivity data and earlier reports (Supplementary Fig. 1A, D-E) (24–27).

### HIV-1 T/F reporter viruses exhibit improved reporter stability

We next asked whether each HIV-1 T/F reporter virus was capable of maintaining stable reporter expression over multiple rounds of replication. We performed this experiment under conditions where fresh uninfected primary CD4^+^ T cells were added every two days to maintain the pressure for the virus to spread to uninfected cells. Primary CD4^+^ T cells were infected with each HIV-1 T/F reporter virus, and after every 48 hours, the percent p24^+^ cells in each culture was determined and maintained at the same level by adding fresh autologous CD4^+^ T cells (between 0.1-1%, depending on the experiment). Under these conditions, CD4^+^ T cells infected with the NL4-3 derived IRES.GFP reporter virus (hereafter called NL43.R5.GFP) used in previous reports (28, 29), quickly lost reporter expression with only about ∼5% of the p24^+^ cells still expressing GFP after 1 week (Fig. 2A-C). In contrast, reporter expression for the HIV-1 T/F reporter viruses remained stable after multiple rounds of replication during the same timeframe (Fig. 2B-C).

**Figure 2.**
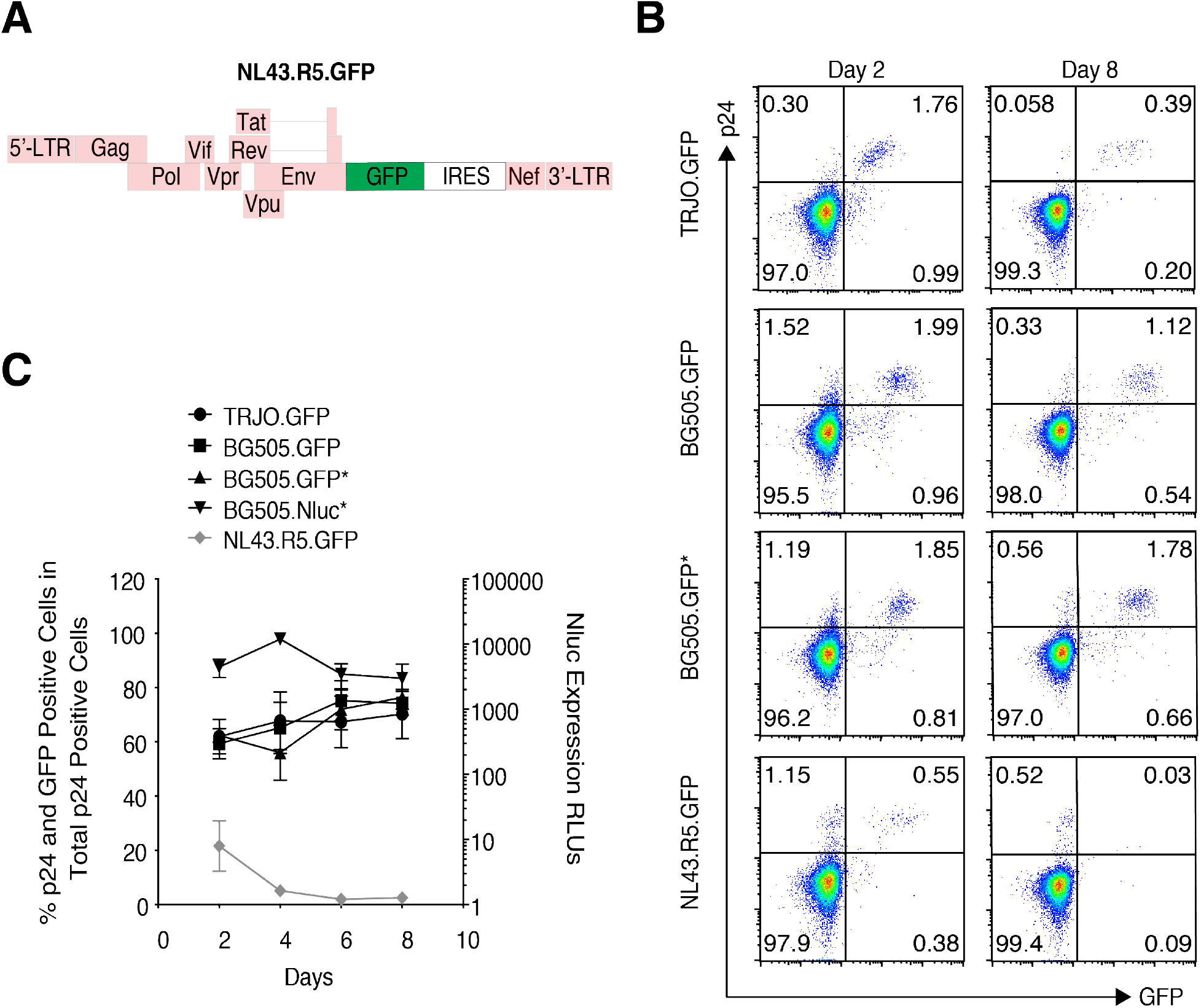
In vitro HIV-1 T/F reporter virus reporter gene stability in primary CD4+ T cells. (A) NL43.R5.GFP reporter virus design. Purple regions correspond to duplicated 3’-LTR sequence flanking the reporter gene. (B, C) Stability of GFP and Nluc reporter viruses in primary CD4 T cells. To force the reporter viruses to continuously spread to new cells, fresh autologous uninfected T cells were added every 48 hours so that the percent of p24+ cells in the total culture was maintained at the same value for each experiment. HIV-1 T/F and NL43.R5.GFP reporter virus GFP and p24 expression 2 days and 8 days post-initiation (B). FACS data is representative of 3-6 independent experiments. Reporter expression was determined via flow cytometry and displayed as the percentage of double-positive HIV-1 and GFP co-expressing cells in the total p24+ population for GFP expressing reporter viruses and the total fold Nluc-derived light units above an Efiverenz negative control for the BG505.Nluc* reporter virus (C). Data in (C) shown as the mean +/-SEM of 3-6 independent donor primary CD4+ T cell preparations.

We next monitored reporter stability of HIV-1 T/F reporter viruses during spreading infection in humanized mice. We selected the GFP expressing BG505.GFP* and BG505.GFP as representatives of the two T/F reporter virus strains used for these studies in order to further investigate the effect of each IRES element on reporter gene stability in vivo. Humanized PBMC mice (Hu-PBL) were infected intraperitoneally (i.p.) with 1 x 10^7^ Infectious Units (IUs) of either BG505.GFP* or BG505.GFP and the percent of GFP expressing cells in the total peripheral blood p24^+^ CD3^+^ T cell population was assessed at different time points over a two-week period via p24 intracellular flow cytometry. We observed that GFP expression in BG505.GFP* infected Hu-PBL mice was very stable for 14 days (Supplementary Fig. 2A). Approximately 30-80% of p24^+^ cells expressed GFP by Day 4 post-infection (Supplementary Fig. 2A-B), and this value increased to approximately 80-90% of p24^+^ cells expressing GFP by Day 11 (Supplementary Fig. 2B). In contrast, GFP expression in BG505.GFP infected Hu-PBL mice was less stable. GFP expressing cells averaged roughly 20% in the total p24^+^ population (Supplementary Fig. 2C). GFP expression persisted for 9 days post-infection and then declined by day 12 post-infection likely due to the observed toxicity of Nef over-expressing cells (Supplementary Fig. 2C). These experiments identify the BG505.GFP* virus as a well-behaved reporter virus.

### GFP expressing reporter viruses productively label infected CD4^+^ T cells in Hu-PBL mice for confocal and two-photon microscopy

To detect GFP reporter expression in parallel, we analyzed spleen tissue in Hu-PBL mice at 7 days post-infection with 1 x 10^7^ IUs of TRJO.GFP. Cryosections of infected spleen tissue was prepared for confocal imaging of GFP and CD3 co-expressing cells. Many infected cells were morphologically distinct and exhibited elongated membrane extensions in comparison to adjacent non-infected cells, suggesting that multinucleated syncytia may have been formed during active infection of compartmentalized lymphocytes in secondary lymphoid tissues (Supplementary Fig. 3A, 1-2 white arrows). We also monitored the behavior of GFP^+^ infected cells within the spleen using intravital laser scanning multiphoton microscopy (LS-MPM) imaging. We infected Hu-PBL mice i.p. with 1 x 10^7^ IUs of TRJO.GFP and adoptively transferred RFP-expressing autologous primary CD4^+^ T cells into infected Hu-PBL mice 24 hours prior to imaging. We observed motile clusters of GFP expressing cells in the spleen with very elongated membrane protrusions (Supplementary Fig. 3B-C) for the full-length T/F virus TRJO.c, reproducing earlier findings (28, 30, 31). We also observed GFP^+^ cells containing multiple foci of concentrated RFP signal, indicative of multiple primary T cells forming a syncytium within 24 hours after exposure to HIV-1 infected cells (Supplementary Fig. 3D-E). In short, TRJO.GFP infected cells exhibit a diversity of cellular morphologies from syncytia to single cells in vivo (Supplementary Fig. 3D-G).

### Nluc enables detection of approximately 30-50 infected cells in vitro and in vivo

We evaluated the sensitivity of Nluc detection in vitro in tissue culture, ex vivo within tissues, and in vivo in living animals. To best correlate infected cells with Nluc expressing cells, we generated a GFP and Nluc coexpressing single-round Gammaretroviral vector from the pMIG MLV-derived vector (hereafter called pMIG.Nluc.IRES.GFP) (Fig. 3A). This vector was designed to express both Nluc and GFP from a single bicistronic transcript to ensure that both reporter gene would be generated in transduced cells at a ratio of approximately 1:1. In addition, we chose to conduct this experiment in wild-type BALB/c mice as they develop native lymphoid organ systems and mature secondary lymphoid organ tissue architecture, in contrast to humanized mouse models, which exhibit a greater amount of variability in lymphoid tissue development and cellular engraftment. Overall, our strategy would provide greater reproducibility and overall accuracy when quantifying Nluc sensitivity estimates under in vivo imaging conditions.

**Figure 3.**
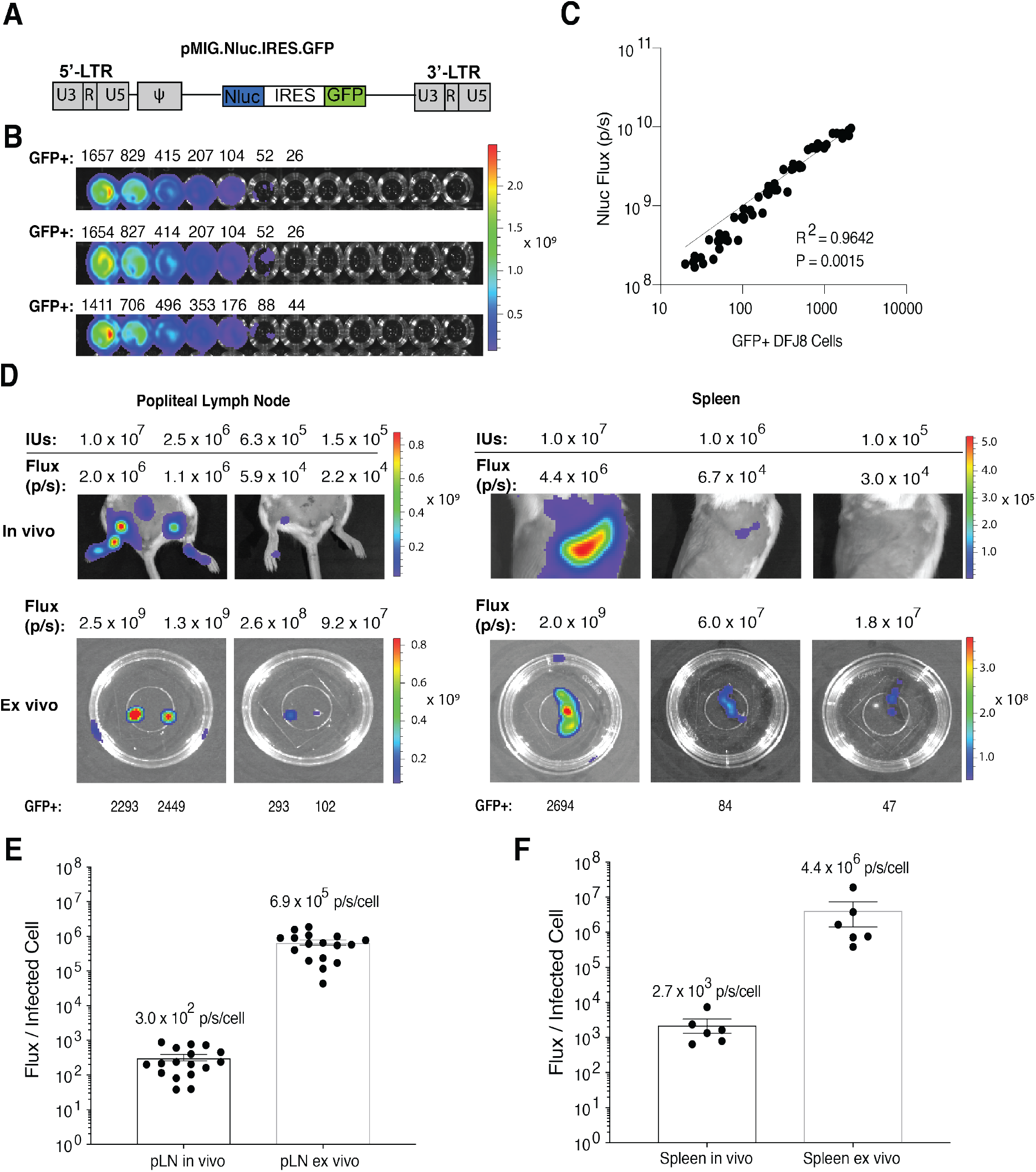
Nluc sensitivity and cellular detection limit in vitro, in vivo and ex vivo. (A) Schematic of the Nluc and GFP co-expressing retroviral vector pMIG.Nluc.IRES.GFP. (B) In vitro Nluc signal sensitivity for Friend-MLV (F-MLV) pseudotyped pMIG.Nluc.IRES.GFP virus-infected DFJ8 cells. DFJ8 cells were infected with 5 x 10^5^ infectious units (IUs) of F-MLV pseudotyped pMIG.Nluc.IRES.GFP virus. Infected cell cultures were resuspended in 150 μl of PBS and then subsequently divided into 50 μl and 100 μl aliquots. The number of infected cells in the 50 μl aliquot (i.e. GFP+ cells) were determined by flow cytometry and the 100 μl aliquot was then serially diluted two-fold on a 96 well plate and imaged using IVIS Spectrum instrument after adding Nano-Glo Nluc substrate at a final ratio of 1:40. Data shown as three representative technical replicates from one of three independent experiments (n=3). (C) Linear regression analysis of GFP expression and Nluc signal flux (i.e. p/s) in infected DFJ8 cells from three independent experiments. (D) In vivo and ex vivo Nluc signal detection limit in BALB/c mice infected subcutaneously via the footpad as well as intravenously via the retro-orbital route with decreasing concentrations of Friend MLV pMIG.Nluc.IRES.GFP virus for delivery to the popliteal lymph node and spleen, respectively. GFP expressing cells were assessed via flow cytometry from cell suspensions generated from tissues following bioluminescent imaging. (E-F) Ratio of total Nluc flux to GFP expressing infected cells for the (D) popliteal lymph node (n=16) and (E) spleen (n=6) imaged both in vivo and ex vivo. Average flux/infected cell ratios are shown above for each imaging condition and each organ. Data shown as mean +/-SEM of 6-16 independent organ preparations.

To determine the Nluc reporter cellular detection limit in vitro, DFJ8 cells were plated in 96 well plates at 1 x 10^6^ cells per well and infected with 5 x 10^5^ IUs of Friend MLV (F-MLV) Envelope pseudotyped pMIG.NLuc.IRES.GFP virus. 48 hours post-infection, the number of GFP^+^ cells was determined by flow cytometry, two-fold serial dilutions were prepared in Nano-Glo Nluc substrate at a dilution of 1:40 in PBS, and samples were then imaged for Nluc bioluminescent signal using an IVIS Spectrum instrument. We observed the Nluc generated flux (photons / second = p/s) dropping to below the detection threshold of the IVIS Spectrum instrument (defined as signal above background levels detected in uninfected controls) when an estimated 30-50 cells were reached (Fig. 3B). Additionally, the relationship between GFP^+^ cells and Nluc signal was strongly linear (Fig. 3C, R^2^ = 0.98, P=0.02), suggesting that Nluc-derived bioluminescence is a good indicator of the level of GFP^+^ cells and can be used to quantify infection levels during longitudinal NIBLI imaging.

We next tested the Nluc sensitivity and Nluc per infected cell ratios in vivo in living animals as well as ex vivo in tissues such as lymph nodes and spleen. BALB/c mice were infected subcutaneously (s.c.) via the footpad or intravenously (i.v.) via the retroorbital (r.o.) route at progressively decreasing infectious doses with single-round F-MLV pseudotyped pMIG.Nluc.IRES.GFP viral particles. Animals were imaged in vivo under anesthesia, and following necropsy, infected tissues from the same animals imaged in vivo were subsequently imaged ex vivo. Cell suspensions were then prepared from infected tissues and the number of infected cells determined by flow cytometry. Under in vivo imaging conditions, we were able to detect between 100-300 cells in popliteal lymph nodes, and between 50-80 cells in the spleen when the mouse was orientated on its lateral side, before the signal became undistinguishable from background levels (Fig. 3D). The spleen is located peripherally beneath the skin, which may explain the higher detection limit observed for this organ. In contrast, the popliteal lymph node is enclosed within adipose and connective tissues and lies behind the muscles of the hind leg, increasing the amount of light diffraction occurring between Nluc-expressing infected cells and the IVIS instrument.

We next asked whether we could use Nluc bioluminescence to quantify cell infection levels in vivo. For each lymph node and spleen analyzed above, we calculated the ratio of whole-tissue Nluc signal flux with the number of GFP^+^ cells determined via flow cytometry (Fig. 3E-F). On average, 3.0 x 10^2^ +/-6.0 x 10^1^ flux units were measured per infected cell in lymph nodes imaged under in vivo conditions and 6.9 x 10^5^ +/-1.2 x 10^5^ flux units per infected cell for lymph nodes imaged under ex vivo conditions (Fig. 3E). Average Nluc-derived flux per infected cell in the spleen was quantified as 2.7 x 10^3^ +/-1.0 x 10^3^ flux units per infected cell measured in vivo and 4.4 x 10^6^ +/-3.0 x 10^6^ flux units per infected cell measured ex vivo (Fig. 3F). These results indicate that under ex vivo imaging conditions, approximately two orders of magnitude more photons can be captured from the same tissue. Considering that the IVIS Spectrum instrument has a dynamic range over about 11 orders of magnitude, our data further supports the use of NIBLI methods to visualize as well as quantify infection levels within a good range of accuracy.

Lastly, we determined the in vitro cellular detection limit of BG505.Nluc* T/F reporter virus infection. TZM-bl indicator cells were infected with 3.4 x 10^5^ IUs of BG505.Nluc* reporter virus in the presence of 16 μg/ml DEAE Dextran. Samples were spinoculated at 1200 x g for two hours at room temperature and allowed to incubate at 37°C for 36 hours. TZM-bl cells from each replicate were then prepared similarly to pMIG.Nluc.IRES.GFP infected DFJ8 described above and subjected to parallel p24 intracellular staining and IVIS imaging (Supplementary Fig. 4). Nluc signal flux dropped below the background detection limit when an estimated 24-50 p24^+^ cells were reached, in close agreement with the 30-50 cell detection limit estimated for pMIG.Nluc.IRES.GFP infected cells (Supplementary Fig. 4A). Additionally, linear regression analysis demonstrated that p24 expression was linearly associated with Nluc expression (R^2^=0.9250, P=<0.0001) (Supplementary Fig. 4B). This data suggests that our original cellular detection limit estimates do not fluctuate greatly between the MLV and HIV-1 LTR promoters driving Nluc expression in our pMIG.Nluc.IRES.GFP and BG505.Nluc* constructs, respectively.

### Q23.BG505.Nluc T/F reporter virus allows for longitudinal NIBLI imaging of HIV-1 dissemination and Nluc signal is strongly correlated with HIV-1 infection

We next performed longitudinal NIBLI imaging of HIV-1 dissemination in Hu-PBL and Hu-HSC mice. Individual Hu-PBL mice (n=6) and Hu-HSC mice (n=2) were infected i.p. with 1 x 10^7^ and 1 x 10^6^ IUs of full-length BG505.Nluc* reporter virus, respectively, and imaged over a period of 15 days. Nluc signal in infected hu-mice exhibited a common spreading pattern under our experimental conditions. By Day 3-6 post-infection, infected cells were first detected in peripheral lymph nodes, such as the mandibular lymph nodes (Fig. 4A-B). From these initial sites, infection continued to spread, reaching the spleen by day 12 and 15 post-infection, for Hu-PBL and Hu-HSC mice respectively (Fig. 4A, B). Whole animal signal flux derived from Nluc expressing cells in infected Hu-PBL mice progressively increased over time with the most rapid increase observed between 3-12 days post-infection (Fig. 4C).

**Figure 4.**
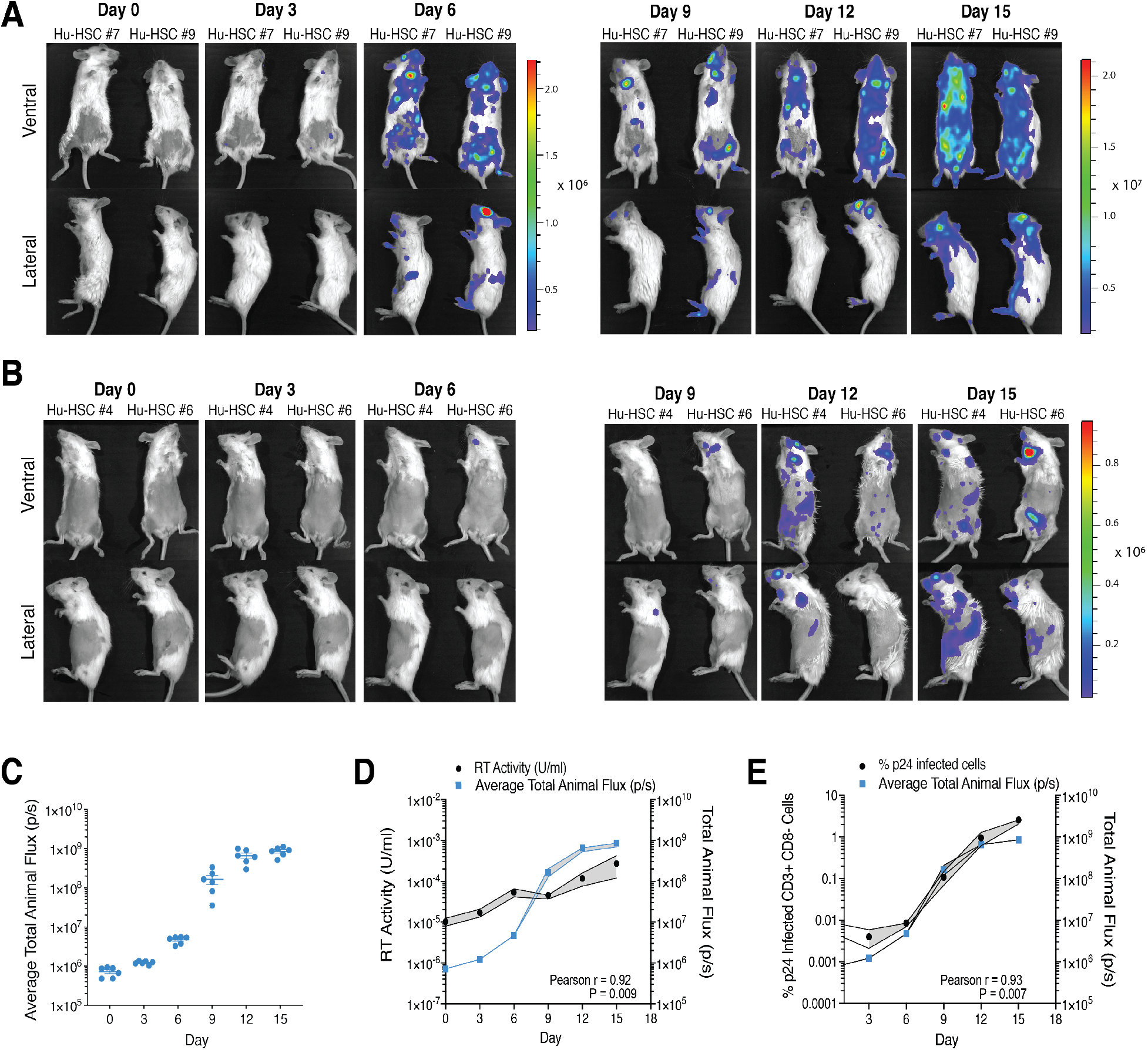
Non-invasive bioluminescent imaging of longitudinal HIV-1_Q23.BG505_ infection in humanized mice. (A) Longitudinal imaging of Hu-PBL mice (n=6) infected intraperitonially (i.p.) with 1 x 10^7^ infectious units (IUs) of HIV-1 BG505.Nluc* reporter virus. Data representative of six independently infected animals. (B) Longitudinal imaging of productive HIV-1 infection in Hu-HSC mice (n=2) infected i.p. with 1 x 10^6^ IUs of BG505.Nluc* T/F reporter virus. (C) Quantification of bioluminescent signal measured over a 15 day period in BG505.Nluc* infected Hu-PBL mice. Average total animal flux was calculated by taking the mean of the total animal flux measured from both the ventral and lateral imaging orientations, excluding the cranial region to avoid signal artifacts arising from the Nano-glo substrate injection site. Data displayed as the mean +/-SEM from six independent experiments. (D, E) Correlative analysis of average total animal flux with the increase of peripheral HIV-1 infected cells (p24+CD3+CD8-cells) measured by flow cytometry (D) and plasma reserve transcriptase activity (RT U) in the infected Hu-PBL mice in (C) measured by the reserve transcriptase SG-PERT activity qPCR assay (E). Upper and lower bounds of the SEM is displayed as gray shaded regions above and below the mean value at each day measured. Pearson correlation calculated from (D) the average values of peripheral HIV-1 infected cells and average total animal flux and (E) plasma reverse transcriptase activity and average total animal flux.

We next wanted to determine how longitudinal Nluc flux measurements related to classic indicators of HIV-1 infection, such as plasma reverse transcriptase (RT) activity and p24 expression. Peripheral blood samples from infected Hu-PBL mice imaged longitudinally via NIBLI above were separated into plasma and PBMC fractions. RT activity was measured on blood plasma using a previously established quantitative PCR (qPCR) based SYBR-Green product enhanced (SG-PERT) RT activity assay (32, 33). Measuring RT activity from virus particles over viral genomes offered the advantage that it would be unaffected by input plasmid DNA contamination present during early time points after infection from virus produced by transient transfection of HEK293 producer cells (32, 33). By removing this possibility, we aimed to provide reliable measurements during very early infection windows. Moreover, the SG-PERT assay was selected due to its high sensitivity and reported detection limit of as low as 3-50 viral particles per reaction (32, 33). We further validated the sensitivity the SG-PERT qPCR assay and compared the assay to a standard qRT-PCR based RNA viral load assay. We performed a series of ten-fold dilutions from a peripheral blood mononuclear cell (PBMC) derived HIV-1 supernatant stock and ran each assay in parallel, using the diluted supernatant as input template. Both assays reached a limit of detection and lost their linear dynamic range between the same dilutions (10^-6^ and 10^-7^) indicating that both assays have comparable sensitivities (Supplementary Fig. 5B). The viral supernatant used for both assays was found to contain 1.63 x 10^9^ RNA copies / mL via the qRT-PCR RNA viral load assay, and considering that the linear range for both assays ended around the 10^-6^ dilution, that an HIV-1 viral particle is known to possess two RNA genome copies, and that the starting input volume for the SG-PERT assay was 15 μL of the same viral supernatant, we estimate that the SG-PERT assay can detect as low as 3-30 viral particles with confidence, very close to previous estimates of 3-50 particles (32, 33).

Applied to HIV-1 spread in hu-mice, the plasma RT activity was strongly correlated with the measured Nluc signal flux (Pearson r = 0.92, P = 0.009) (Fig 4C). Similarly, the increase in average p24^+^ cell levels measured in plasma by flow cytometry was highly correlated with the average whole-animal signal flux during the course of the 15-day infection (Pearson r = 0.93, P = 0.007) (Fig. 4D). Together, these data show that RT activity as measured by the SG-PERT qPCR assay is a suitable method for determining plasma viremia during HIV-1 infection.

### Recrudescent infection is detected in tissues initially infected with HIV-1 in Hu-HSC and Hu-BLT mice

To determine whether we could monitor HIV-1 infection during cART and recrudescent infection after cART interruption, we infected hu-mice i.p. with 1 x 10^6^ IUs of BG505.Nluc* reporter virus (n=5). cART was initiated during early time points post-infection in order to capture the early seeding of a long-lived cellular reservoir established during the onset of acute infection (34). Infection was allowed to progress for 6 days for one group (2 Hu-HSC mice), 9 days for the second group (2 Hu-HSC mice), and 12 days for one Hu-BLT mouse before cART was initiated. The cART regimen consisted of daily i.p. injections of Truvada (TDF/FTC) and Isentress (RAL) at recommended doses for humanized mice (200 mg/kg TDF, 133 mg/kg FTC, and 80 mg/kg RAL) (35).

We imaged longitudinal HIV-1 infection in these mice before, during, and after cART cessation (Fig. 5A, Fig. 6A, Supplementary Fig. 5). Daily cART was administered for 30 days for Hu-HSC mice and 24 days for Hu-BLT mice and interrupted at day 36 post-infection. During the cART treatment period, the average total animal flux gradually faded with a concomitant decrease in the plasma viremia, measured in parallel via the SG-PERT assay (Fig. 5A-B, Fig. 6A-B), and Supplementary Fig. 6A,C). For all mice, plasma viremia dropped to the day 0 uninfected baseline by 6-12 days after the initiation of daily cART injections and remained suppressed until cART withdrawal (Fig. 5B, Fig. 6B, Supplementary Fig. 6C). One of the 4 Hu-HSC mice showed a viremic blip on day 18, but was suppressed throughout the remainder of the cART treatment period (Fig. 6B).

**Figure 5.**
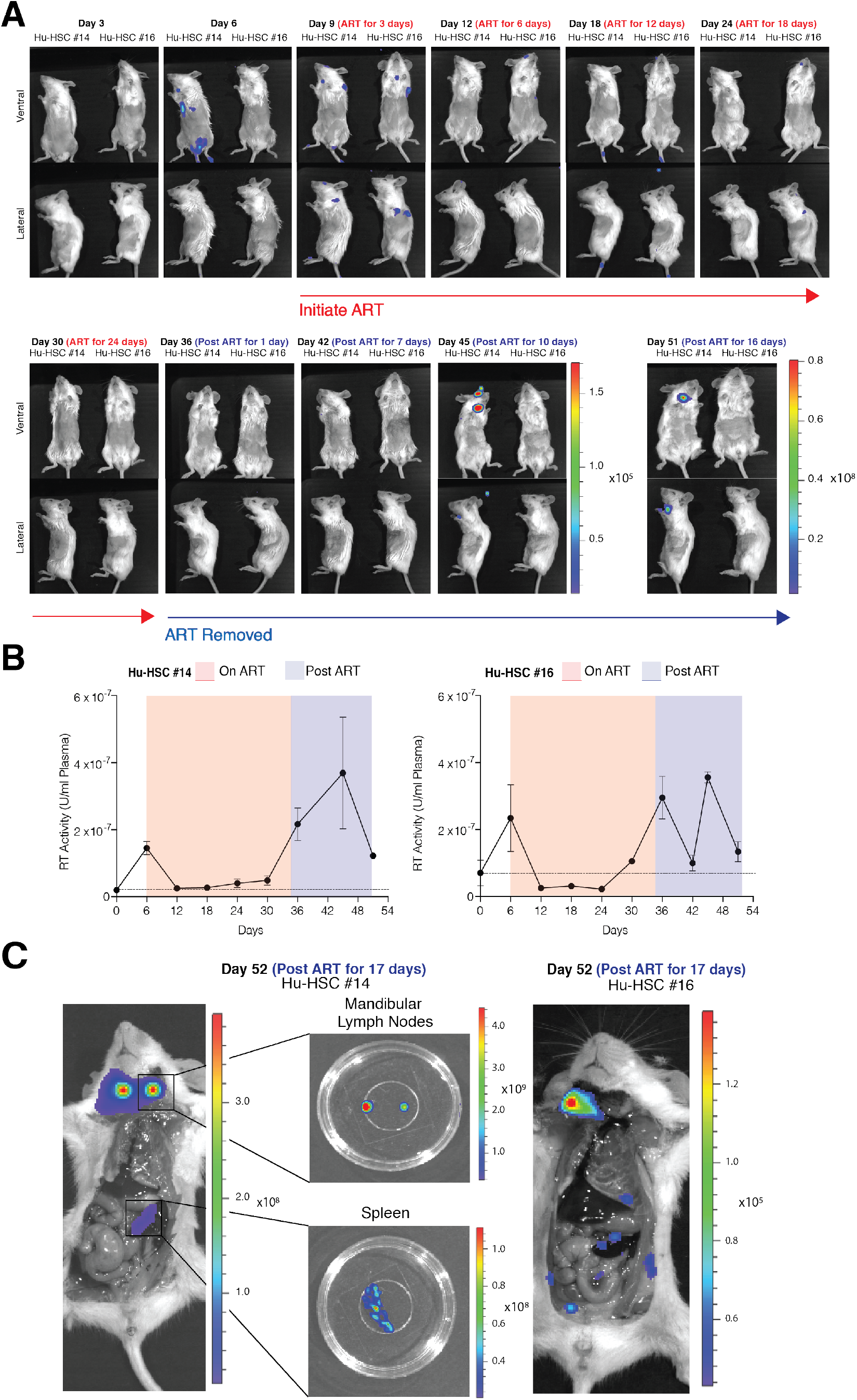
Longitudinal imaging of HIV-1 acute infection, cART suppression, and infection recrudescence in Hu-HSC mice placed on ART 6 days post-infection. (A) Longitudinal bioluminescent imaging of spreading infection in Hu-HSC mice infected with 1 x 10^6^ infectious units (IUs) of BG505.Nluc* T/F reporter virus and placed on a daily ART regimen comprised of daily i.p. ART injections of Truvada and Isentress 6 days post-infection (n=2). (B) Quantification of plasma reverse transcriptase activity from the mice in (A). Plasma reverse transcriptase activity in serum samples obtained every six days over the course of the imaging period was measured via the SG-PERT reverse transcriptase activity assay and described as reverse transcriptase activity units / mL above endogenous uninfected background levels (shown as a dotted line). (C) Whole animal ex vivo necroscopic analysis of recrudescent infection in Hu-HSC mice from (A) approximately two weeks following ART cessation.

**Figure 6.**
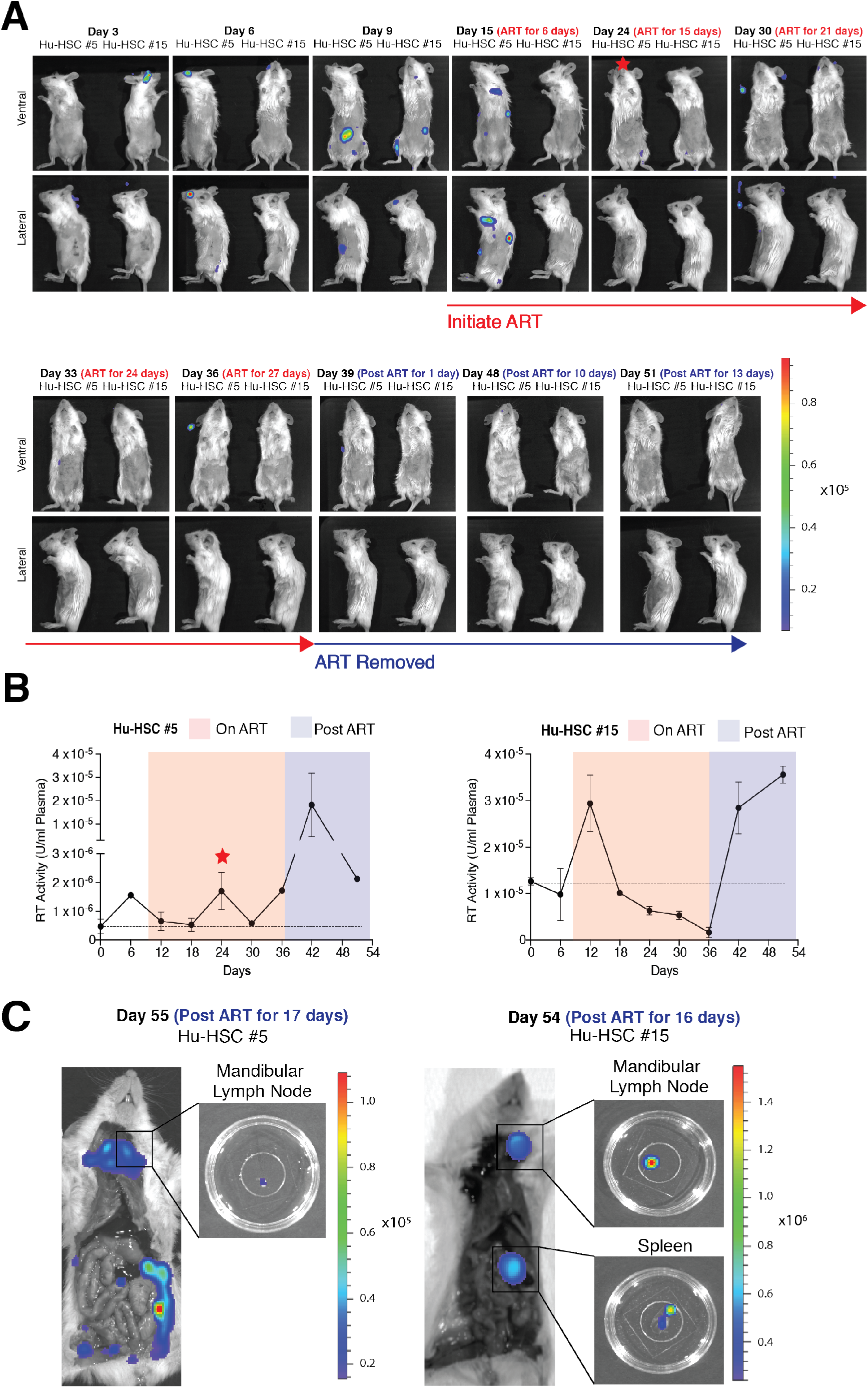
Longitudinal imaging of HIV-1 acute infection, suppression, and recrudescent infection in Hu-HSC mice placed on ART 9 days post-infection. (A) Longitudinal bioluminescent imaging of HIV-1 acute infection, suppression, and recrudescent infection in Hu-HSC mice infected with 1 x 10^6^ IUs of BG505.Nluc* T/F reporter virus and placed on daily i.p. ART injections of Truvada and Isentress 9 days post-infection (n=2). Red star denotes the timepoint and Hu-HSC mouse that exhibited a transient increase in plasma reverse transcriptase activity during ART treatment. (B) Quantification of plasma reverse transcriptase activity from the animals in (A). Plasma reverse transcriptase activity in serum samples taken every six days over the course of the imaging period was measured via the SG-PERT reverse transcriptase activity assay and described as reverse transcriptase activity units / mL above endogenous uninfected background levels (shown as a dotted line). (C) Whole animal ex vivo necroscopic analysis of recrudescent infection in Hu-HSC mice from (A), approximately two weeks following ART cessation.

Recrudescent infection was observed for all mice, either in vivo or following necropsy performed after the termination of the experiment (Fig. 5A,C, Fig. 6A,C, Supplementary Fig. 6A-B). Rebound occurred in the mandibular lymph nodes above the clavicle bones and medial to the sublingual and submandibular salivary glands where the infection was first observed in 4 out of 5 mice (Fig. 5A,C: Hu-HSC mouse #14 and Hu-HSC mouse #16, Fig. 6A,C: Hu-HSC mouse #15, and Supplementary Fig. 6A-B: Hu-BLT Mouse #3). Necroscopic analysis of each animal revealed rebound infection in the spleen in two of these animals (Fig. 5C: Hu-HSC mouse #14 and Fig. 6C: Hu-HSC mouse #15). Notably, we observed an increase in plasma viremia one time point prior to the IVIS signal in the tissue due in part to the higher sensitivity of the SG-PERT assay for some of the mice analyzed (Fig. 5A-B: Hu-HSC Mouse #16 and Fig. 6A-B: Hu-HSC Mouse #5). Interestingly, comparative analysis of Nluc expression before cART initiation and after cART interruption showed that for all mice analyzed, secondary lymphoid tissues positive for Nluc signal during the recrudescent infection often harbored HIV-1 infected cells before the initiation of cART (Fig. 5A, C, Fig. 6A, C, Supplementary Fig. 6A, B). These data suggest that a cellular reservoir may be established early in these mice (before 6 days), in agreement with observations made in SIV-infected NHP studies (34).

### HIV-1 infected T cells and macrophages were detected in Nluc expressing tissues during recrudescent infection

We isolated tissues positive for Nluc signal during infection recrudescence to characterize HIV-1 infected tissues at the single-cell level. Hu-HSC mouse tissues that exhibited positive endogenous Nluc signal following cART treatment interruption were surgically removed, fixed in 4% paraformaldehyde in PBS and subsequently cleared for cryoimmunofluorescent confocal microscopy (Fig. 7A). In addition to identifying p24^+^ CD3^+^ lymphocytes, as expected, we observed infection of CD68^+^ macrophages (Fig. 7 B-C). HIV-1 infected CD68^+^ macrophages were observed in multiple lymphoid tissues in at least two Hu-HSC mice (Fig. 7C, Hu-HSC Mouse #14 and Hu-HSC Mouse #15, data not shown).

**Figure 7:**
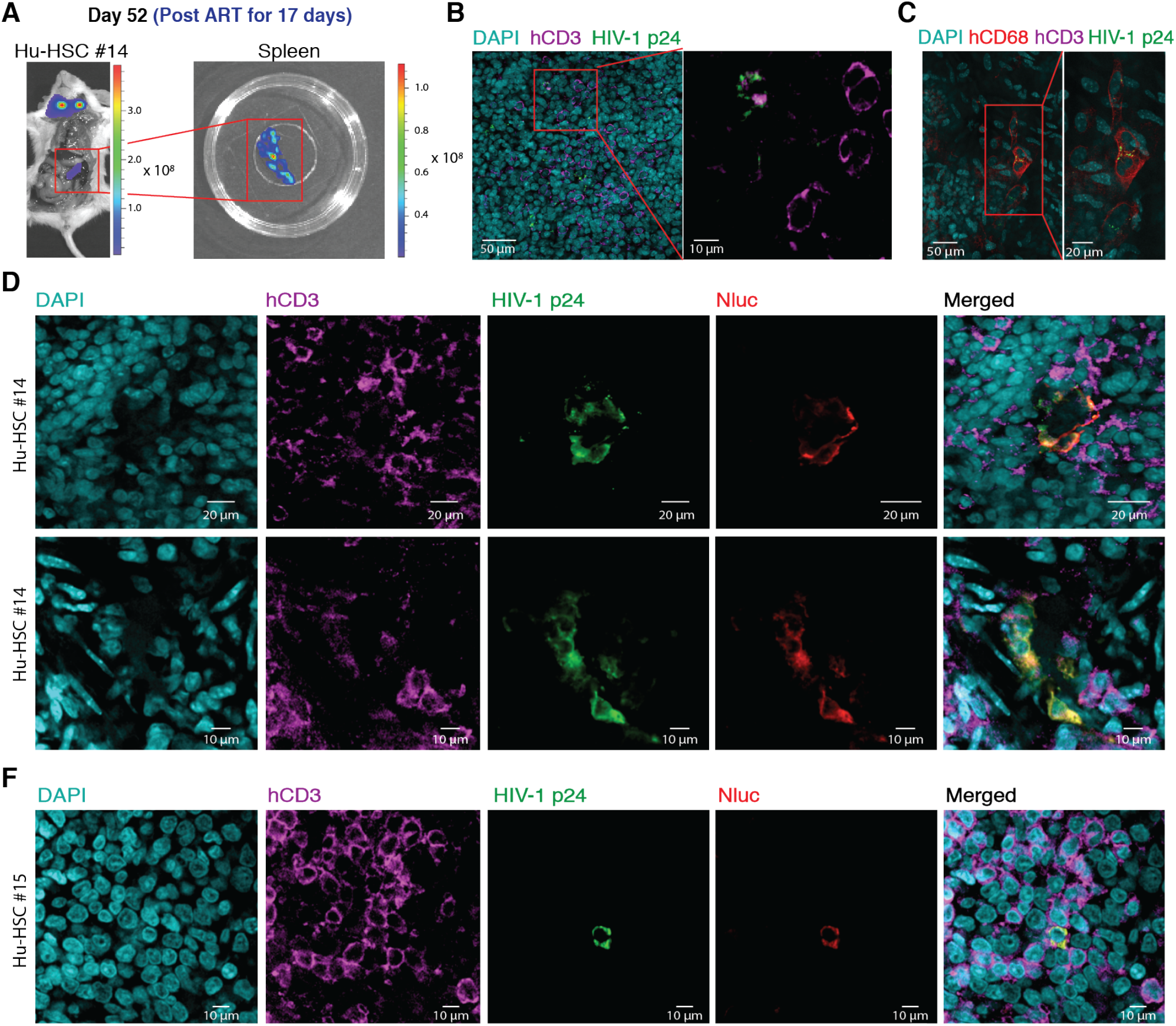
Confocal immunofluorescence microscopy of cleared HIV-1 Q23.BG505.Nluc infected spleen tissue during recrudescent infection. (A) Nluc expressing spleen tissue from Hu-HSC mouse #14 infected with BG505.Nluc* reporter virus. Nluc expressing spleen is enclosed in red boxes at whole-animal and tissue resolutions, respectively. Spleen tissue was surgically removed 17 days following cART withdrawal and was subsequently fixed, cleared, and immunostained for confocal microscopy to identify HIV-1 p24+ cells. (B-C) Representative confocal slices of Nluc expressing spleen tissue from (A). Tissues were immunostained for HIV-1 p24 (green), human CD3+ T cells (magenta), human CD68+ macrophages (red), and stained with DAPI to identify nuclei (cyan). HIV-1 p24 was associated with human CD3+ T-cells (B) and human CD68+ macrophages (C) within the same piece of Nluc expressing spleen tissue. (D-F) Immunostaining of representative Nluc expressing cells in spleen tissue from BG505.Nluc* infected Hu-HSC mouse #14 (D) and Hu-HSC mouse #15 (F). Nluc expression (red) colocalizes with HIV-1 p24 (green) and CD3 (magenta) in spleen tissue isolated from both Hu-HSC mice.

To further test whether the Nluc signal was indeed derived from HIV-1 infected cells during recrudescent infection, we immunostained additional tissues from BG505.Nluc* infected Hu-HSC mice isolated approximately two weeks following cART withdrawal with a monoclonal antibody against Nluc (Fig. 7 D-F). We found that Nluc specifically colocalized with HIV-1 p24 in spleen tissue isolated from two different Hu-HSC mice (Fig.7 D, Hu-HSC mouse #14 and Fig. 7 F, Hu-HSC mouse #15). By confirming the presence of Nluc in cells positive for p24 in tissues, we demonstrated that the Nluc signal detected during whole-animal and ex vivo tissue imaging was derived from cells that were HIV-1 infected. Taken together, these results demonstrate the utility of Nluc-based longitudinal NIBLI to identify HIV-1 productive infection during different disease and treatment conditions at whole-animal, tissue, and cellular resolutions.

## Discussion

In this report, we have applied NIBLI to visualize longitudinal HIV-1 viral dissemination, suppression under cART, and infection recrudescence following treatment withdrawal. This was possible due to the generation of replication-competent full-length reporter viruses from transmitted / founder (T/F) HIV-1 strains expressing GFP or Nluc upstream of the *nef* open reading frame. By inserting an internal ribosome entry site (IRES) sequence between the reporter gene and *nef*, Nef expression and function was preserved to near-native levels. In vivo, between 30-50 infected cells in superficial tissues and 100-300 infected cells in collagen-rich tissues such as lymph nodes could be detected via Nluc bioluminescence. In addition, Nluc expression was strongly correlated with total infected cells and reverse transcriptase activity in blood plasma and the signal gradually decreased when mice were treated with daily cART. Longitudinal NIBLI of humanized mice following cART withdrawal allowed us to localize tissue sites involved in viral recrudescence. Interestingly, we observed that infection rebound was observed in tissue sites that were positive for active HIV-1 replication prior to ART initiation.

Our results suggest that HIV-1 recrudescent infection emerges from tissue-sites that harbored infection during the earliest stages of acute infection. The presence of Nluc signal during infection rebound, under conditions in which cART was initiated very early during acute infection, further suggests that a long-lived cellular reservoir harboring proviral sequences derived from BG505.Nluc* infection is rapidly established (34, 36). Notably, the absence of Nluc expression in most lymphoid tissues following cART withdrawal suggests a fairly static cellular reservoir during the cART suppression period. This observation lends credence to the possibility that a subset of the long-lived cellular reservoir established during HIV-1 infection in humanized mice has tissue-resident characteristics.

We also demonstrated how whole-body NIBLI can direct subsequent higher resolution imaging to infected tissues and identify HIV-1 infected cells by confocal imaging. Interestingly, this approach identified CD3^+^ T cells as well as CD68^+^ macrophages in splenic tissues. The role of macrophages as potential reservoirs for HIV-1 is still unclear. In vitro, macrophages have been observed to be infected by HIV-1 directly, by phagocytosis of HIV-1 infected T cells, or by syncytia formation with HIV-1 infected T cells (41–44). Whether macrophage reservoirs critically contribute to long-term persistence of HIV-1 following cART treatment interruption in patients requires further scrutiny.

The high replicative capacity and rapid replication time of HIV-1 exerts a strong selection pressure to maintain genomic integrity and remove any introduced gene that does not confer a fitness advantage. Our results showed that we were able to increase the stability of our reporter viruses over previous variants by preserving the native genome structure in conjunction with using reporter genes smaller than 1 kilobases (kb), which includes GFP (∼800 bp) and Nluc (∼500 bp). This stability was also dependent on whether the full-length ECMV IRES or the truncated and attenuated 6ATRi variant was used to drive *nef* expression. Overall, our T/F reporter constructs possessing the 6ATRi IRES element exhibited superior reporter gene stability and better resembled infection by wild-type virus. These findings underscore the need to regulate HIV-1 accessory gene levels to avoid unwanted cytotoxicity and other ectopic effects from reporter viral infection.

Being mindful about the strengths and limits of these reporter viruses, we maintain that they can be useful tools for *in vivo* studies particularly in small animal models such as humanized mice. As humanized mouse models that better recapitulate human immune responses against infectious diseases become available, we are hopeful that the HIV-1 T/F reporter virus toolkit described herein can make important contributions to ongoing HIV-1 research and cure development programs.

## Materials and Methods

### Ethics Statement

All animal experiments were performed according to protocols approved by the Institutional Review Board and the Institutional Animal Care and Use Committee (IACUC) of Yale University. Yale University is registered as a research facility with the United States Department of Agriculture (USDA), License and Registration number: 16-R-0001. It also is fully accredited by the Association for Assessment and Accreditation of Laboratory Animal Care (AAALAC). An Animal Welfare Assurance (#A3230-01) is on file with OLAW-NIH.

### Construction and preparation of HIV-1 T/F GFP and Nluc reporter viruses

GFP and Nluc HIV-1 reporter viruses were assembled using Gibson Assembly, modified from the original published method (45). In short, assembly was performed using equimolar amounts of the following PCR amplified fragments: GFP and Nluc open reading frames, using the pHXB.eGFP.IRES expression vector (a gift from Heinrich Gottlinger, Worcester, UMass, MA) and pNL.1.1 from Promega, respectively; IRES and 6ATRi sequences from the PHXB.eGFP.IRES vector and the pNL43.GFP.6ATRi.Nef infectious molecular clone (IMC) (a gift from Dr. Christina Oschenbauer), respectively; and adjacent regions of the pTRJO.c and Q23.BG505 IMCs, using primers with 40 base pair overlaps. PCR fragments were mixed 1:4 with a mixed enzyme assembly master mix cocktail, and incubated at 50°C for 30 minutes to 1 hour. The assembled insert was then cloned into wildtype pTRJO.c or Q23.BG505. Validated HIV-1 T/F reporter virus plasmid was transfected into HEK293 cells (ATCC, Manassas, VA) and cultured at 37°C in RPMI supplemented with 10% fetal calf serum (FCS) and 1% penicillin / streptomycin (i.e. RPMI cell culture medium) (Thermo-Fischer Scientific, Waltham, MA) in a 1:3 DNA volume / transfection reagent volume ratio with Fugene 6 transfection reagent (Promega, Madison, WI), using the standard manufacturer’s instructions. Culture supernatant was saved and replaced every 24 hours for 48 hours, filtered through 0.45 μM syringe filters (Pall Corporation, Port Washington, NY) and stored at −80°C. Virus preparations were titered on TZM-bl cells (NIH AIDS Reagents) seeded at a density of 1 x 10^5^ cells per well in a 24-well tissue culture dish (Corning) in 400 μL RPMI cell culture medium containing 16 μg / ml DEAE-Dextran (Sigma, St. Louis, MO). Plates were spinoculated at 1200 x g for 2 hours at room temperature and incubated at 37°C for 48 hours. Medium was replaced with 200 μL Accutase Solution (Stemcell Technologies, Vancouver, Canada) and incubated for 5 to 10 minutes at 37°C. Cells were then washed and fixed in 100-200 μL 1X Cytofix / Cytoperm solution (BD Biosciences, Franklin Lakes, NJ) and incubated a 4°C for 20 minutes. Cells were then resuspended in 100-200 μL 1X Perm / Wash Buffer (BD Biosciences) containing anti-HIV-1 p24-RD2 (Beckman Coulter, clone KC57) diluted 1:1000, and incubated at 4°C for 30 minutes. Cells were then resuspended in PBS + 2% BSA and analyzed via flow cytometry using a BD Accuri C6 flow cytometer to quantify p24 positive cells. Titers in infectious units (IUs) were then calculated as (((1 x 10^5^) x (% p24 positive cells) x (dilution factor)) / (volume in mL))) to give IU / ml, as previously published (46).

### Single round infectivity assays

TZM-bl cells were plated at 1 x 10^5^ cells / 100 μL RPMI + 10% fetal calf serum / well into 48 well plates and infected with 100 μL viral supernatant with 16 μg / ml DEAE-Dextran and 10 nM of the protease inhibitor saquinavir. Plates were spinoculated for 1200 x g for 2 hours at room temperature and infection was allowed to proceed for 48 hours at 37°C for 48 hours. Culture media was then removed and cells were resuspended in 200 μL firefly luciferase passive lysis buffer (Promega). Lysate was then placed into white-bottom 96 well plates (Corning) at 20 μL per well. Luciferase readings were performed by dispensing 25 μL of firefly luciferase substrate (Promega) and measuring the subsequent bioluminescent output via a TriStar LB 941 multimode reader (Berthold). Relative light units were then normalized to an efavirenz uninfected control and then to reverse transcriptase units / ml viral supernatant.

### Western blotting for Nef protein expression

2.25 x 10^6^ HEK293 cells were transfected with 15 μg of each HIV-1 virus plasmid DNA in T75 flasks (for TRJO.GFP western blots) and 5 x 10^5^ HEK293 cells were transfected with 2 μg HIV-1 plasmid DNA in six well plates (for BG505.GFP, BG505.GFP*, and BG505.Nluc* western blots), each using Fugene 6 (Promega) at a 1:3 DNA:Fugene 6 ratio. Cells in each well were collected and washed in PBS, spun at 8000 rpm for 5 minutes, and resuspended in 150 μL chilled RIPA Buffer (Boston Bioproducts, Ashland, MA) and incubated for 10 minutes on ice. Lysates were collected after spinning each sample at 14,000 rpm for 30 minutes at 4°C and removing the supernatant into a fresh 1.5 ml Eppendorf tube containing 50 μL of 4X NuPAGE protein sample buffer (Life Technologies) (950 μL 4X LDS Buffer + 50 μL 1M DTT). Samples were heated at 80°C in a heat block for 5 minutes with shaking and stored at −20°C if not used immediately. Samples were then run in 4-12% NuPAGE SDS-PAGE protein gels (Life Technologies) submerged in 1X MOPS Buffer (Boston Bioproducts) at 150 V until dye front had migrated off the gel. Gels were then placed above 0.2 μm Nitrocellulose filters between two pieces of Whatman filter paper (Bio-Rad Laboratories, Hercules, CA) and transferred in 10X Transfer Buffer (0.4M Tris + 2M Glycine in dH_2_O) diluted to 1X Transfer Buffer with a 7:2 vol / vol ratio of dH_2_O and methanol for 2 hours at 2 Amps. The membrane was washed 4 times with cold dH_2_O and incubated with primary antibody (anti-HIV-1 Nef antiserum or anti-HIV-1 p24 clone 183-H12-5C, both from NIH AIDS Reagents) diluted 1:1000 in PBST / BSA blocking buffer (PBS + 0.05% Tween 20 + 5% BSA) overnight at 4°C on a rocker. Membranes were then washed 3 times with PBST buffer and incubated with secondary antibody (either conjugated with Horseradish Peroxidase, HRP, or anti-rabbit / anti-mouse Fc Alexa-680 / Alexa-780) diluted 1:15000 in PBST + 5% powdered milk for 1 hour rocking at room temperature. Membranes were then washed 3 times with PBST. Chemiluminescent signal was detected via an ImageQuant LAS 4000 (GE, Fairfield, CT) and fluorescent signal via LiCor Odyssey Classic (LiCor, Lincoln, NE) using the manufacturer’s software.

### Nef functionality assays

Nef functionality was determined by seeding 1 x 10^6^ JLTRG-R5 cells (NIH AIDS Reagents) per well into a 96 U bottom plate (Corning) at a final volume of 50 μl. 2 ml of frozen HIV T/F and HIV T/F reporter virus stocks were defrosted at 37°C for 1 minute and concentrated using Lenti-X concentrator, following the manufacturer’s instructions into 50 μl working stocks using RPMI + 10% FCS + 1% Penicillin / Streptomycin +1% glutamate cell culture medium with 16 μg / ml DEAE-Dextran added. Efavirenz controls were included by adding efavirenz to a final volume of 1 μM into the cell culture medium and incubated at 37°C for 1 hour prior to the addition of virus. Virus was then dispensed into each well and spinoculated at 1200 x g for 2 hours at room temperature and allowed to incubate at 37°C for 48 hours. Each sample was assays for intracellular p24 via flow cytometry, as previously described above, and all samples were then equilibrated to the lowest p24 positive population (0.6%) by adding fresh JLTRGFP.R5 cells (NIH AIDS Reagents). Infection was then allowed to proceed for 48 hours and CD4 levels were determined by measuring the mean fluorescent intensity of CD4 surface expression via flow cytometry using anti-CD4 conjugated to Alexa 647 (BioLegend, San Diego, CA clone OKT4). CD4 expression was compared between GFP expressing (i.e. HIV-1 infected) and the total CD4 positive population from the uninfected efavirenz control.

### Primary cell reporter stability assays

PBMCs were isolated from leukopacks (New York Blood Center), diluted with room temperature PBS 1:5 and added to 50 ml falcon tubes above a 14 ml layer of Ficoll Plaque Plus (GE Health Care, Chicago, IL). Tubes were spun at 1200 x g for 30 minutes at room temperature and the central leukocyte fraction was carefully removed and washed 3 times in room temperature PBS to remove traces of Ficoll. PBMCs were then resuspended in RPMI + 10% FCS / 1% Penicillin / Streptomycin and CD4 T cells were magnetically isolated using a negative selection CD4 T cell immunoisolation kit (Stemcell Technologies) following the manufacturer’s instructions. Freshly isolated primary cells were then resuspended in RPMI + 10% FCS / 1% Penicillin / Streptomycin with 100 ng/ml human IL-2 (PeproTech, Rocky Hill, NJ), 10 ng/ml human IL-7 (R&D Systems, Minneapolis, MN), and 10 ng/ml IL-15 (R&D Systems) and cultured in T25 filtered cull culture flasks (Eppendorf, Hamburg, Germany). For spreading assays, primary T cells were plated in 96 U bottom plates at 1 x 10^5^ – 1 x 10^6^ cells / well depending on the experiment. After virus was introduced, plates were spinoculated at 1200 x g for 2 hours at room temperature and allowed to incubate at 37°C for 48 hours. A sample of cells was analyzed for intracellular p24 levels by flow cytometry and fresh autologous primary T cells were then added to each well to equilibrate all wells to the lowest p24 level detected (0.1 – 1 % depending on the experiment). Fresh primary T cells were added every 48 hours for 8-12 continuous days keeping p24 levels at the same percentage of total p24 positive cells throughout the experiment.

### In vitro HIV-1 T/F reporter virus p24 and GFP coexpression analysis

For each experiment, 1 x 10^5^ CEM.SS.R5.NKr cells were suspended in 50 μl cell culture medium (RPMI + 10% FCS + 1% Penicillin / Streptomycin + 1% Glutamate) and 16 μg / ml DEAE Dextran with each sample plated in triplicate in U bottom 96 well plates. HIV-1 T/F reporter virus supernatant was concentrated with Lenti-X concentrator, using the manufacturer’s protocol, and resuspended in cell culture medium containing 16 μg / ml DEAE Dextran at a concentration of 1 x 10^5^ IUs / 50 μl. RT controls contained efavirenz at a final concentration of 1μM. Virus was added to each sample for a MOI of 1 and a final volume of 100 μl. Each plate was spinoculated at 1200 x g for 2 hours at room temperature and immediately placed into a cell culture incubator at 37°C / 5% CO_2_. Samples were prepared for p24 intracellular flow cytometry at the specified time point using the Cytofix/Cytoperm buffer protocol with monoclonal mouse anti-HIV-1 p24-RD2 (KC57, Beckman Coulter) at a 1:1000 dilution.

### Generation of humanized mice

Hu-PBL and Hu-HSC hu-mice were generated from the xenoengraftment of human hematopoietic cells into immunocompromised NOD.*Cg-Prkdc^scid^Il2rg^tm1Wjl^*/SzJ (NSG) mice (Jackson Laboratories, Bar Harbor, ME), maintained under the daily care of the Yale Animal Resources Center (YARC). For generating Hu-HSC mice, 1-2 day old neonatal NSG mice (Jackson Laboratories, Bar Harbor, Maine) were irradiated at 100 cGy and injected through the intracardiac route with 1 x 10^5^ CD34^+^ hematopoietic stem cells purified from human umbilical cord blood (National Disease Research Interchange). Hu-PBL mice were generated by i.p. injection of 1 x 10^7^ PBMCs stored on ice into 2-4 month old NSG mice. Mice were analyzed via flow cytometry after 10 weeks for Hu-HSC or after 2 weeks for Hu-PBL for the successful engraftment of monoclonal mouse anti-human CD45 (BioLegend, clone HI30), monoclonal mouse anti-human CD3 (BioLegend, clone HIT3a), monoclonal mouse anti-human CD4 (BioLegend, clone RPA-T4), and monocloncal mouse anti-human CD8 (BioLegend, clone RPA-T8) positive cells until all mice exhibited greater than 25% human CD45 expressing cells.

### In vivo reporter stability in Hu-PBL mice

Hu-PBL hu-mice expressing a minimum of 25% CD45^+^ cells were i.p. injected with approximately 1 x 10^7^ IU of T/F reporter virus. Infections were allowed to proceed for 16 days and mice were bled retro-orbitally at 3 days post infection. 100 μl of blood was diluted 1:1 with fresh sterile PBS and overlaied above 100ul of lymphoprep (Stemcell Technologies) for PBMC extraction. Samples were centrifuged for 20 minutes at 2000 rpm in a bucket router and the PBMC fraction was isolated and stained with monocloncal mouse anti-human CD3 (BioLegend, clone UCHT1) and monoclonal mouse anti-HIV-1 p24-RD2 (Bekman Coulter, clone KC57) at 1:100 and 1:1000 dilutions, respectively using the Cytofix/Cytoperm kit and accompanying protocol (BD Biosciences). Flow cytometry was performed on a BD C6 Accuri and analyzed with Flowjo vr. 10.

### In vitro and in vivo assessment of Nanoluciferase sensitivity

The Nluc open reading frame was cloned from plasmid pNL.1.1 (Promega) using PCR with forward primer 5’-ATTATCTCGAGCACCATGGTCTTCACA-3’ and reverse primer 5’-TACATGAATTCCCTTACGCCAGAATGC-3’ and was introduced upstream of the IRES element in vector pMIG (Addgene plasmid #9044) via restriction digest with XhoI and EcoRI to generate pMIG.Nluc.IRES.GFP. For in vitro assessments, DFJ8 cells were plated in triplicate in 96 U bottom plates at a concentration of 1 x 10^6^ cells / well and infected in the presence of 16 μg/ml DEAE Dextran with 5 x 10^5^ IUs of retroviral vector pMIG.Nluc.IRES.GFP decorated with ecotropic Friend Murine Leukemia virus envelope, spinoculated at 1200 x g for 2h, and incubated at 37C for 48 hours. Cells were then resuspended in 150 μl PBS and distributed into 50 μl and 100 μl aliquots. GFP expressing cells in the 50 μl aliquot were determined via flow cytometry and 12 two-fold dilutions were prepared in PBS from the 100 μl aliquot. NanoGlo substrate (Promega) was then added at a final ratio of 1:40 in PBS. Plates were then imaged using an IVIS Spectrum Instrument (Perkin Elmer) adjusted to auto-exposure imaging settings. For in vivo assessments, F-MLV pseudotyped pMIG.Nluc.IRES.GFP was injected either subcutaneously or retro-orbitally in wild-type BALB/c mice at indicated IUs and infection was allowed to proceed for 48 hours. Whole animals were then imaged using the IVIS Spectrum via auto-exposure settings following r.o. injection of 200 μL Nano-Glo substrate diluted 1:40 in PBS. Subsequently, whole tissues were surgically removed and submerged in diluted Nano-Glo substrate (1:40) before imaging. Whole lymph nodes and spleens were then submerged in RPMI media + 50 mM HEPES without serum, treated with collagenase for 20 min at 37 C and then passed through a 70 μm nylon cell strainer (Corning) to obtain single cell suspensions. Splenocytes were additionally treated with Red Blood Cell Lysis Buffer (Sigma). Cell suspensions were then washed with PBS + 2 mM EDTA and resuspended in 500 μL MACS Buffer (PBS + 2% BSA + 1 mM EDTA) to be placed in a 5 ml round bottom polystyrene tube with a cell straining cap (Falcon), spun at 800 rpm for 5 minutes, and resuspended in 700 μL of PBS + 2% BSA. Total GFP expressing cells per lymph node or spleen were then determined via flow cytometry. To test the Nluc expressing cell detection limit of cells infected with BG505.Nluc*, 1 x 10^5^ TZM-bl cells were plated in triplicate in a 96 U bottom plate in the presence of 16 μg/ml DEAE Dextran. Each sample were infected with approximately 3.4 x 10^4^ IUs of BG505.Nluc* and plates were then spinoculated at 1200 x g for 2 hours at room temperature. Infection was allowed to proceed for 48 hours at which point TZM-bl cells were serially diluted two-fold and analyzed by p24 intracellular flow cytometry and IVIS imaging in parallel, similarly to pMIG.Nluc.IRES.GFP infected DFJ8 cells described above.

### Intravital LS-MPM spleen imaging

Humanized Hu-PBL mice intended for multiphoton laser-scanning microscopy of HIV-1 infected spleen tissues were infected i.p. with approximately 1 x 10^7^ IU of TRJO.GFP.IRES.Nef T/F reporter virus and infection was allowed to proceed for 7 days. To image CD4 T cells, the pAPM lentiviral vector was digested with XbaI and MluI and dsRed and dtTomato were cloned between the SFFV promoter and the WPRE element. Mice were injected i.p. with 1 x 10^6^ primary CD4 T cells transduced with a mixture of pAPM.DsRed and pAPM.dtTomato lentiviruses pseudotyped with VGV-G at an MOI of 10 and tested via flow cytometry to ensure >10% CD4 T cells were expressing the reporter. Mice were anesthetized by i.p. injection of a mixture of Ketamine / Xylazine at a final concentration ratio of 15 : 1 mg/ml and immobilized on a custom built stage, which included a 37°C heat pad. The spleen was surgically exposed and placed onto a custom-built stage warmed to 37°C, immersed in PBS and sealed with a coverslip. Mice were kept anesthetized via the delivery of nebulized Isoflurane / O_2_ at a constant flow rate. Multiphoton images and movies were taken with an Olympus BX51WI fluorescence microscope with a 20 x 1.00 NA water immersion Zeiss objective and single-beam LaVision TriM Scope II (LaVision Biotec) using Imspector software. The microscope was customized with a Chameleon Vision II Ti:Sapphire laser (Coherent) with pulse precompensation. Emission wavelengths were collected with photomultiplier tubes (Hamamatsu, Hamamatsu City, Japan) 567-647 nm (RFP) and 500-550 nm (GFP). For 3D still images, 22 optical Z sections were taken 3 μm apart using an assigned wavelength of 900 nm and ascending laser power from 8 to 11%. For 4D movies, 27 optical Z sections were taken 3 μm apart, with an assigned wavelength of 900 nm and an ascending power of 8 to 11.5% for 1 hour.

### Immunostaining of cryosections and confocal microscopy

For histological analysis of TRJO.GFP infected Hu-PBL spleens, tissue samples reserved for histological sectioning were fixed in PBS containing 4% paraformaldehyde (PFA) (Electron Microscopy Sciences, Hatfield, PA) overnight at 4°C under gentle rocking. Samples were washed in PBS and equilibrated with 30% sucrose in PBS for 12 hours at 4°C under gentle rocking. Tissue samples were then submerged in O.C.T medium (Sakura Tissue Tek, Torrance, CA) and then stored at −80°C. Tissue molds were then cut into 20-30 μm sections using a CM1950 Cryostat (Leica, Wetzlar, Germany) and quickly placed onto Superfrost Plus glass microscope slides (Fisherbrand, Pittsburgh, PA) stored at −20°C until stained. Slides were then defrosted to room temperature and washed with PBS to remove the O.C.T. layer. Exposed tissue was then blocked for 1 hour at room temperature with blocking buffer (PBS + 2% BSA) and stained with primary antibody diluted 1:333 with blocking buffer overnight at 4°C. Primary antibody staining solution was removed and slides were washed 5 times with washing buffer (PBS + 0.1% Triton X-100) and mounted with Prolong Diamond mounting medium (Life Technologies) and with glass cover slips. Images were acquired with an TCS SP5 Confocal Microscope System (Leica) using the LAS-X software platform and analyzed via Volocity image analysis software. The primary antibody used in this study was monoclonal mouse anti-human CD3-alexa 647 (BioLegend, clone UCHT1).

For histological analysis of Nluc expressing spleen tissue from BG505.Nluc* infected Hu-HSC mice, Nluc expressing spleen samples were cleared, immunostained, and imaged as previously described (47). Briefly, tissues were excised and fixed overnight in ice-cold 0.1 M Sodium Cacodylate trihydrate (Sigma) buffer containing 8% paraformaldehyde (Electron Microscopy Sciences) and 5% sucrose (Sigma). Fixed tissues were washed in PBS and cleared using the CUBIC protocol (48). Samples were placed in CUBIC-1 buffer (25% w/v urea (Sigma), 25% w/v *N,N,N’,N’*-tetrakis (2-hydroxypropyl) ethylenediamine (Sigma), 15% v/v polyethylene glycol mono-*p*-isooctylphenyl ether/Triton X-100 (Sigma) in PBS), and incubated at 37°C with slight rocking until splenic tissue was decolorized. Samples were then washed in PBS and placed in CUBIC-2 buffer (50% w/v sucrose, 25% w/v urea, 10% w/v 2,2,2’-nitrilotriethanol (Sigma), and 0.1% v/v Triton X-100 (Sigma) in PBS), and incubated at 37°C with gentle rocking until tissue was transparent. Tissues were then washed in PBS and immunostained or stored in PBS at room temperature.

Cleared spleen samples were cut into ∼0.5mm thick sections with a surgical scalpel and blocked in 4% v/v fetal bovine serum (Hyclone), 2% v/v rat anti-mouse CD16/32 (Biolegend), 0.1 % v/v Triton X-100 (Sigma), and 0.01% w/v Sodium Azide (Sigma) in PBS. Samples were stained with primary antibodies (1:200) in blocking buffer without rat anti-mouse CD16/32 for 3 days at room temperature with gentle rocking. Tissues were washed in wash buffer (0.1% v/v Triton X-100 (Sigma) and 0.01% w/v sodium azide (Sigma) in PBS) and incubated with secondary antibodies (1:1000) in blocking buffer without rat anti-mouse CD16/32 for 3 days at room temperature with gentle rocking. Tissues were washed in wash buffer and nuclei were stained with 1 µg/mL 4’,6-diamidino-2-phenylindole (DAPI) in PBS for 10 minutes. Samples were washed and refractive indexed matched in CUBIC-2 buffer overnight prior to mounting and imaging. Primary antibodies used in this experiment were polyclonal rabbit anti-human CD3 (Dako), monoclonal mouse anti-human CD68 (Dako, Clone PG-M1), polyclonal goat anti-p24 (Creative Diagnostics), and monoclonal mouse anti-Nano-Luc (R&D Systems). Conjugated secondary antibodies used in this study were Alexa fluor 647 donkey anti-goat IgG, Alexafluor 488 donkey anti-rabbit IgG, and Alexafluor 594 donkey anti-mouse IgG.

Refractive index matched spleen samples were mounted in CUBIC-2 buffer between two No.0 coverslips (Electron Microscopy Sciences) adhered to 0.5 mm silicone isolators (Electron Microscopy Sciences). Tissues were imaged on a Zeiss LSM-800 with a LD LCI Plan-Apochromat 25 × 0.8 NA Imm Corr DIC M27 multi-immersion objective (w.d. 0.57 mm). The Fiji software suite (Schindelin, et al., 2012) was used to process confocal images. Individual confocal images were smoothed to reduce background autofluorescence.

### Longitudinal bioluminescent imaging of hu-mice and correlative analysis of Nluc flux with p24 expression and RT activity in Hu-PBL mice

Hu-HSC mice were infected i.p. with approximately 1 x 10^6^ IUs of BG505.Nluc* full length HIV-1 reporter virus. At various time points, mice were anesthetized with nebulized Isoflurane / O_2_ in a air-sealed chamber, and then injected intravenously via the r.o. route with 100 μl Nanoglo substrate (Promega) diluted 1:40 in sterile PBS. For experiments involving cART, cART treatment consisted of daily i.p. injections of a combination of Truvada (TDF/FTC) and Isentress (RAL), dissolved in sterile water to achieve a final concentration of 200 mg/kg TDF, 133 mg/kg FTC, and 80 mg/kg RAL (ART was a gift from Dr. Richard Sutton, Yale School of Medicine Department of Internal Medicine). Plasma viremia were determined by measuring RT activity in plasma, isolated from 100 μl of blood drawn retro-orbitally and separated using lymphoprep (Stemcell Technologies) to separate plasma and leukocyte fractions from blood, via the SG-PERT assay, as previously described (32, 33). Necroscopic analysis was performed by surgically exposing internal tissue layers and overlaying fresh Nano-glo substrate diluted 1:40 in sterile PBS onto exposed tissues. Images were taken with an IVIS Spectrum (Perkin-Elmer) under auto-exposure settings and analyzed with the manufacturer’s Living Image software package.

For p24 and RT activity correlative analysis, 100 μL of blood was drawn via retro-orbital eye bleeds and plasma and PBMC fractions were isolated via lymphoprep, as described above, from infected Hu-PBL mice imaged longitudinally. Intracellular p24 staining was performed on extracted PBMCs and p24 expressing cells were quantified using flow cytometry, as described previously, and RT activity was determined via the SG-PERT assay.

### Comparative RNA viral load assay and RT activity assay sensitivity assessment

Total viral RNA was extracted from 140 μL of PBMC derived HIV-1 JR-CSF viral supernatant via the QIAamp Viral RNA Mini Kit (Qiagen) following the manufacturer’s protocol. Total viral RNA was eluted into 60 μL and diluted 1:20 in sterile PBS and 10 fold serial dilutions were performed. RNA copies for each sample were determined simultaneously using the QuantiTect SYBR Green RT-PCR kit (Qiagen) as described above, and absolute RNA copies were calculated from a standard curve run in parallel using HIV-1 89.6 genomic RNA quantified at 2 x 10^8^ copies / mL. Reverse transcriptase activity was measured via the SG-PERT assay as described above from 10 μL of the same viral supernatant diluted 1:20 in sterile PBS and assay lysis buffer (as described in (32)). Subsequent 10 fold dilutions were performed as was done for RNA copy number assessment. All samples were run simultaneously and concurrently with an recombinant HIV-1 reverse transcriptase standard curve (Worthington Biochemical Corporation, Lakewood, NJ).

### Statistics

All descriptive statistics, one-way ANOVA to determine significance in CD4 surface down-modulation experiments to test Nef functionality, and Pearson correlation calculations to determine an association between p24^+^ cells and bioluminescent signal during longitudinal imaging of Hu-PBL mice were performed using GraphPad Prism software, version 7.

## Acknowledgements

We thank Dr. Jonathan Grover, and Dr. Maolin Lu for critical reading of the manuscript, Dr. Thomas Mooruka and Dr. Thorsten Mempel for reagents, Dr. Xaver Sewald for his assistance with LS-MPM intravital spleen imaging, Ruoxi Pi for assistance with flow cytometry analysis, Dr. Richard Sutton for providing us with Truvada and Isentress, Dr. Sebastian Hannemann for training on and access to the LiCor Odyssey Classic Imaging platform, The Yale Center for Science and Social Science Information (CSSSI) StatLab Consultants for support with statistical analyses, and the Yale Center for Cellular and Molecular Imaging for training on the Leico SP8 Confocal Microscope and Volocity Imaging Software.

---

**Supplementary Figure 1:**
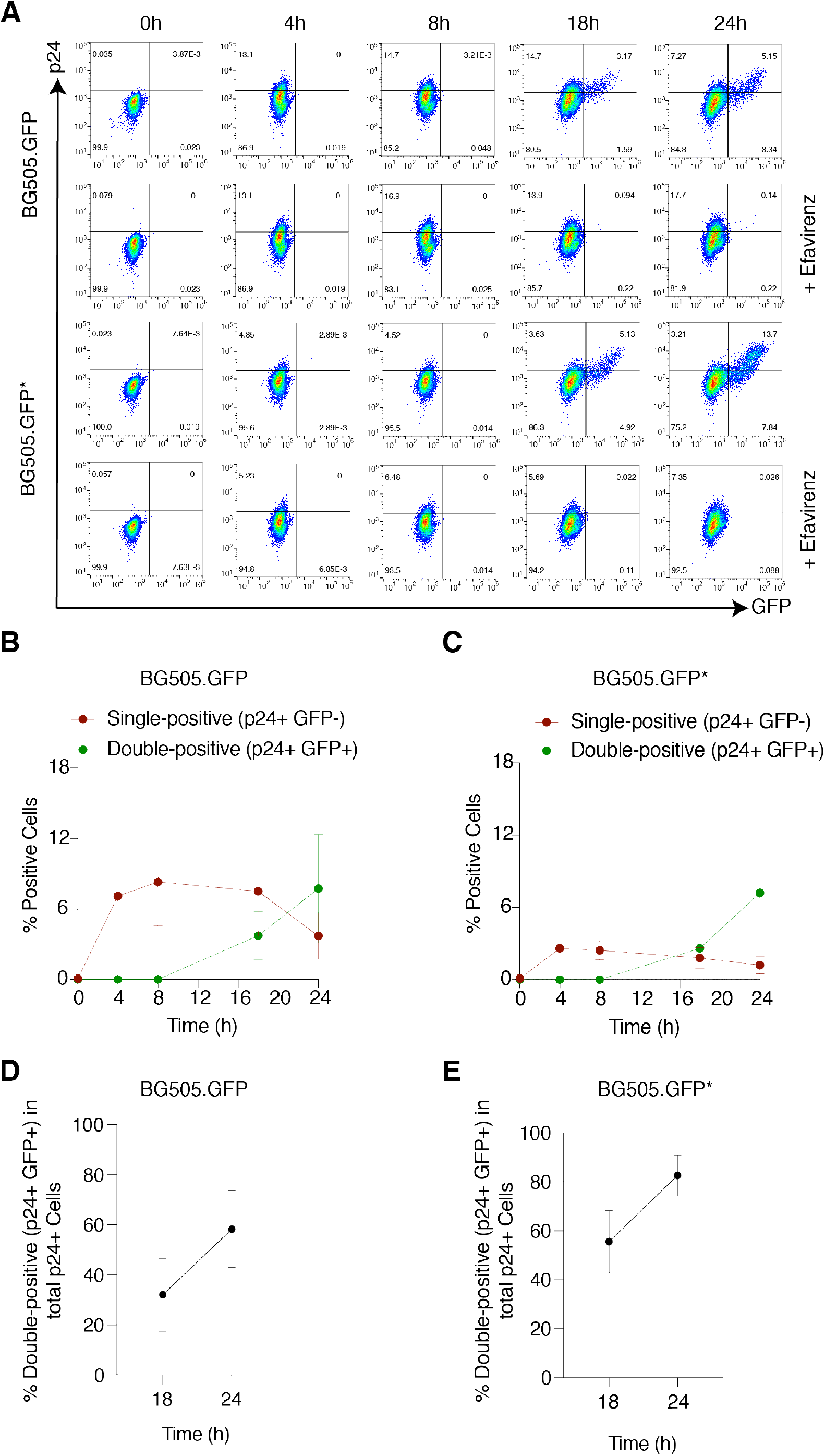
Q23.BG505 T/F reporter virus reporter gene expression kinetics in infected CEMSS.NKr.R5 cells. (A) GFP reporter gene expression at different times points during the initial 24 hours of infection in CEMSS.NKr.R5 cells with or without the addition of 1 μM Efavirenz. Data is representative of three independent experiments. (B-C) Percent of single positive (p24+ GFP-) and double-positive (p24+ GFP+) cells over 24 hours in CEMSS.NKr.R5 cells infected with (B) BG505.GFP or (C) BG505.GFP* T/F reporter viruses. Data displayed as the mean +/-SEM of three independent experiments (n=3). (D-E) The fraction of double-positive cells in the total HIV-1 infected (p24+) population 18 hours and 24 hours post-infection in CEMSS.NKr.R5 cells infected with (D) BG505.GFP or (E) BG505.GFP* T/F reporter viruses.

**Supplementary Figure 2:**
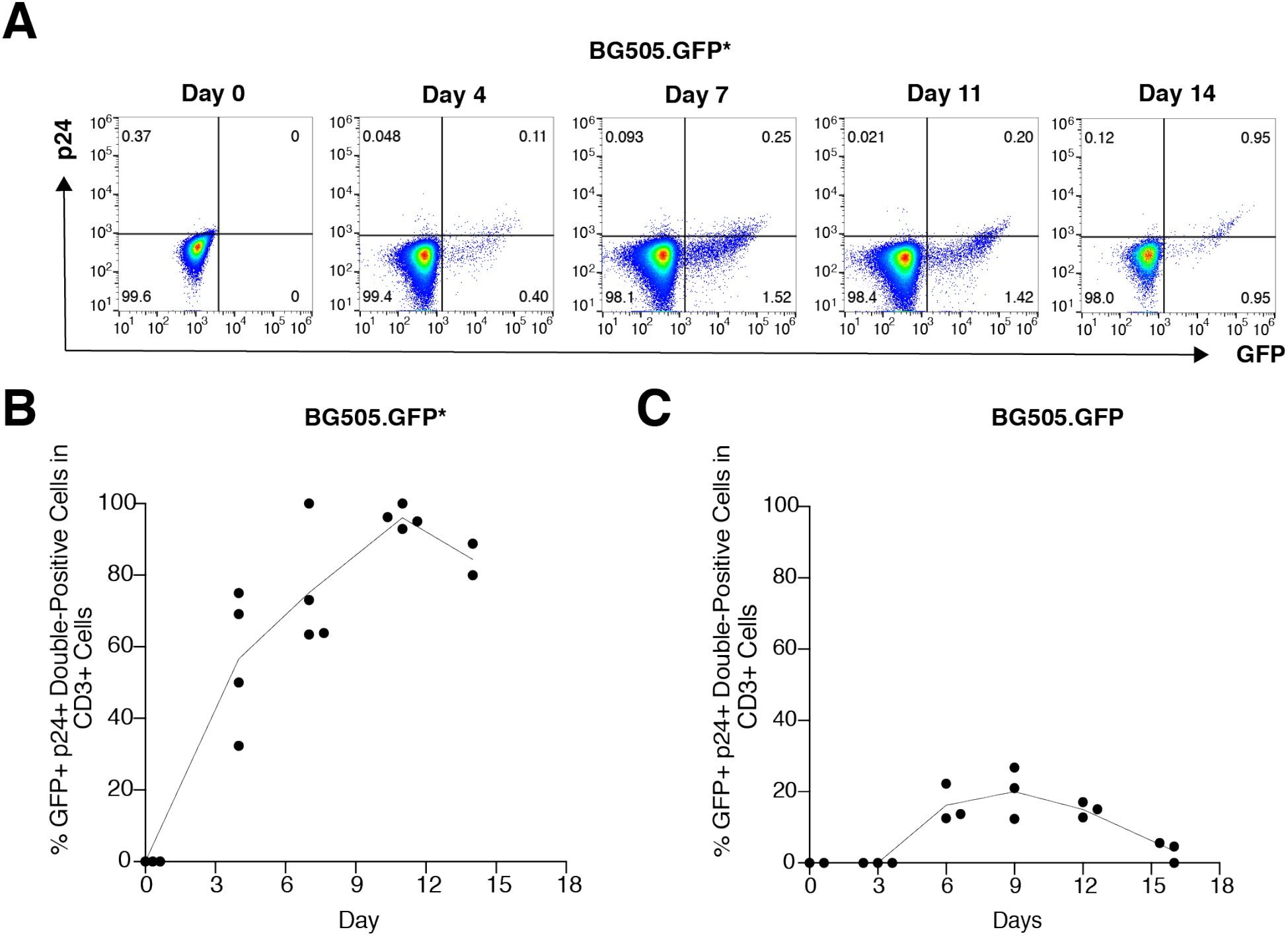
In vivo reporter gene stability in Hu-PBL mice. (A-C) GFP reporter gene stability in p24+ CD3+ PBMCs extracted from the peripheral blood of Hu-PBL mice infected intraperitonially (i.p.) with 1 x 10^7^ infectious units (IUs) HIV-1 T/F reporter virus. (A) Representative FACS plots gated on CD3+ PBMCs extracted from BG505.GFP* infected Hu-PBL mice (n=4). Data is representative of four individual Hu-PBL mice. (B-C) Average GFP reporter gene stability in Hu-PBL mice infected with 1 x 10^7^ infectious units (IUs) of BG505.GFP* (n=4) (B), and BG505.GFP (n=4) (C) T/F reporter virus for 14-16 days. Data displayed as the percentage of GFP and p24 double-positive cells in the total p24+ population. A line crosses the average percent GFP expressing cells within the total p24 + cell population for mice analyzed at each time point.

**Supplementary Figure 3:**
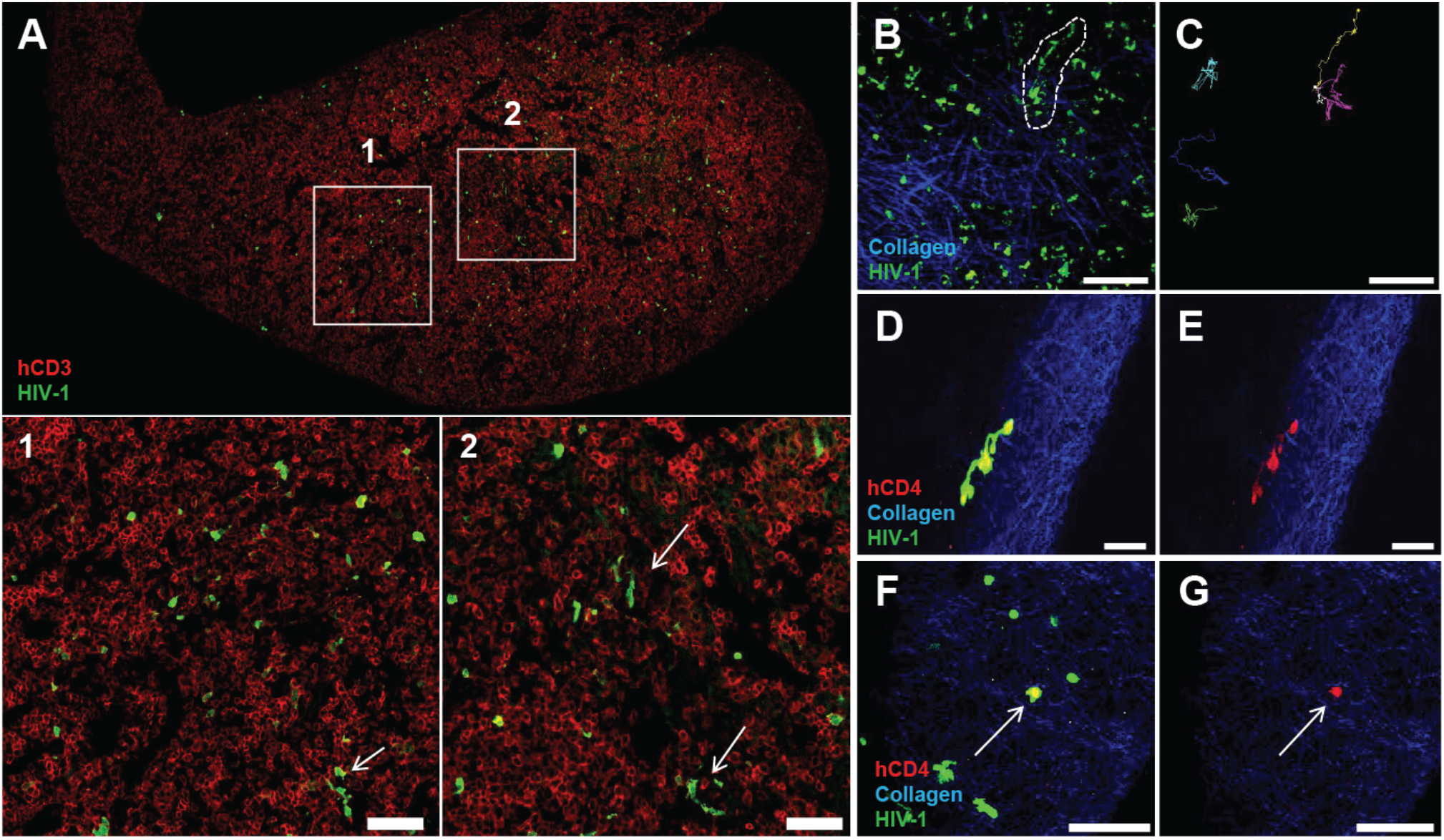
Cryoimmunofluroescent and LS-MPM intravital spleen imaging of Hu-PBL mice injected i.p. with 1 x 10^7^ IUs TRJO.GFP 7 days post-infection. (A) Cryoimmunofluorescent confocal imaging of splenic tissue sections; areas with GFP expressing cells are magnified in panels 1 and 2. White arrows indicate putative syncytia formed during infection. (B-G) LS-MPM imaging of spleen tissue from a Hu-PBL mouse injected i.p. with 1 x 10^7^ IUs TRJO.GFP 7 days post-infection and injected with RFP expressing CD4 T cells 24 hour prior to imaging. (B,C) LS-MPM intravital imaging of an area in the spleen with GFP expressing cells. A representative cell exhibiting long membrane extensions is outlined in white dashes (B) with motion tracks of GFP expressing cells in (C). (D-E) LS-MPM image of GFP and CD4 co-expressing syncytium in the spleen of a TRJO.GFP-infected Hu-PBL mouse (D) and the same image with CD4 expression alone (E). (F-G) LS-MPM image of GFP expressing cells in the spleen as in (D) with a GFP and CD4 co-expressing cell indicated by the white arrow and CD4 expressing cells alone (G). All scale bars correspond to 100 μm.

**Supplementary Figure 4:**
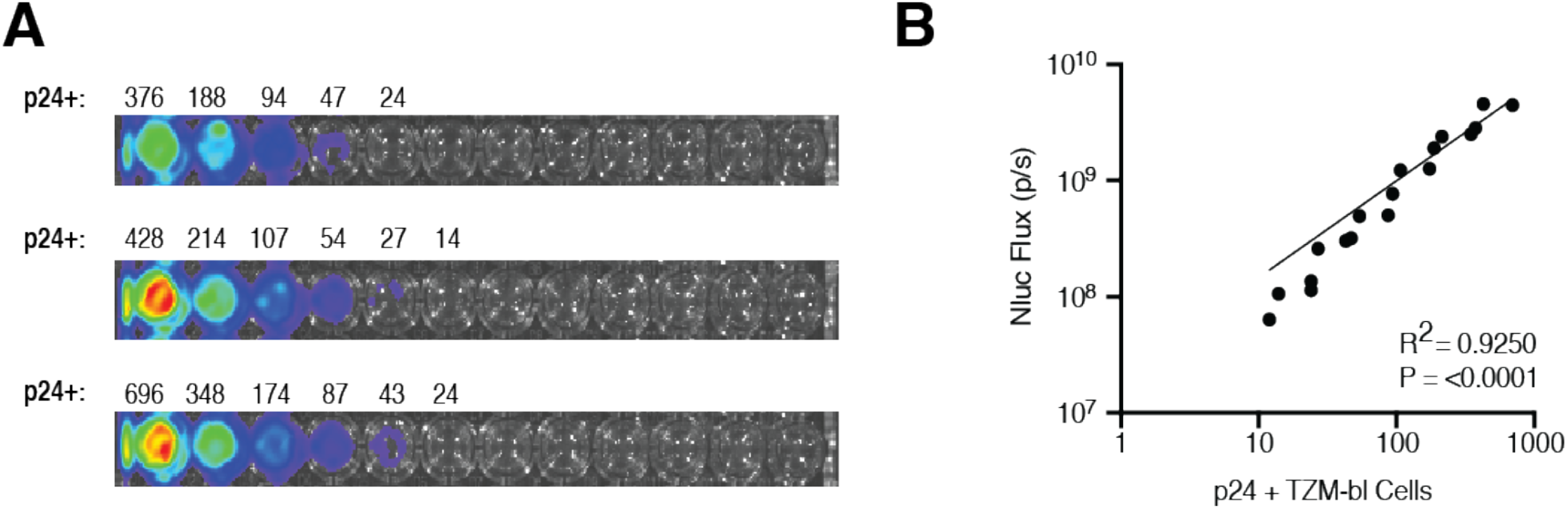
In vitro Nluc signal sensitivity for BG505.Nluc* infected TZM-bl cells. (A) TZM-bl cells were infected with 3.4 x 10^4^ infectious units (IUs) of BG505.Nluc* T/F reporter virus. After 48 hours, each sample was prepared for p24 intracellular flow cytometry and then distributed in two-fold serial dilutions for IVIS imaging. The number of p24 positive cells detected by flow cytometry is placed above the corresponding well analyzed via IVIS imaging. Data shown as three technical replicates from one experiment. (B) Linear regression analysis of p24 expressing cells and total Nluc signal flux (i.e. p/s) from one independent experiment.

**Supplementary Figure 5:**
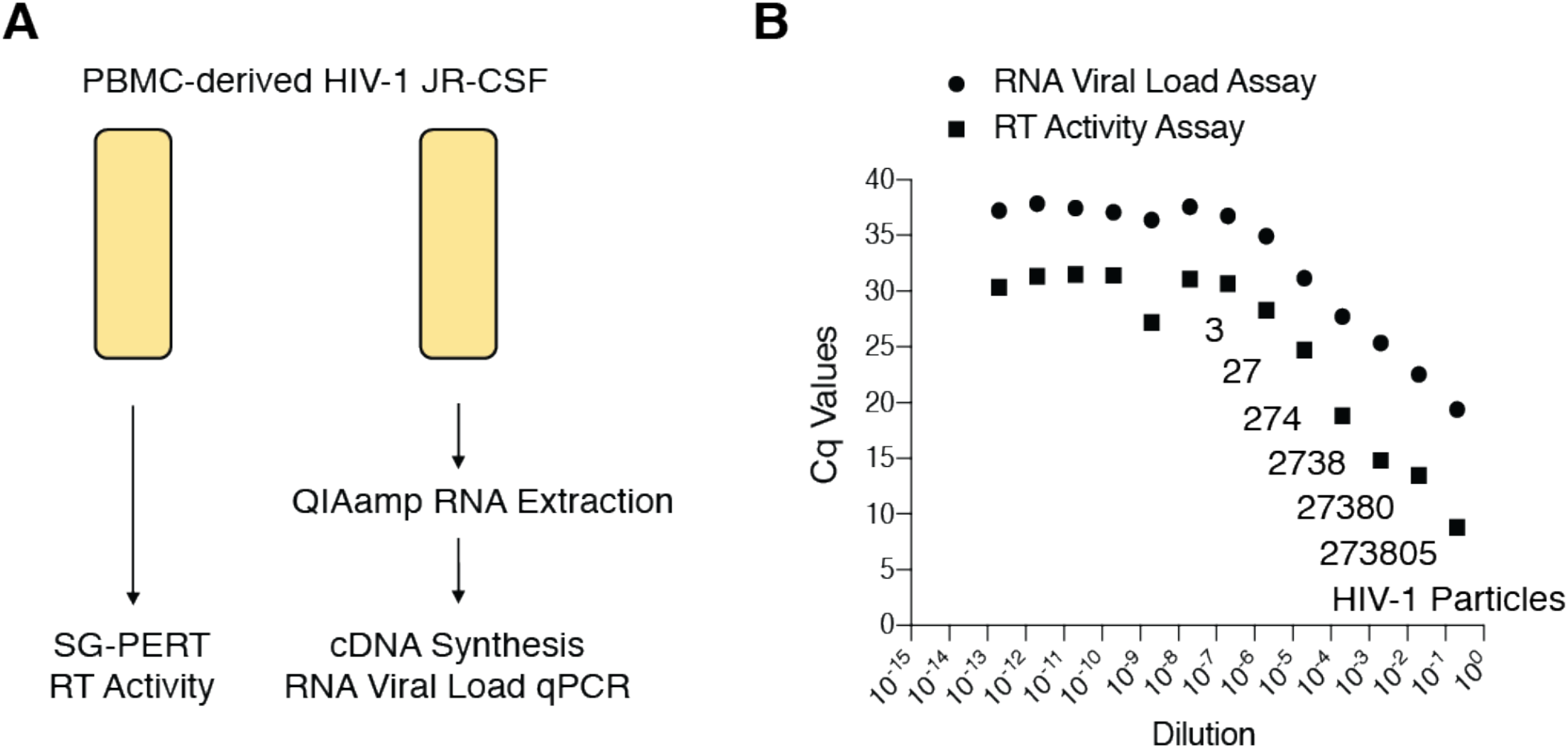
RNA viral load assay and SG-PERT RT activity assay sensitivities. (A) eripheral blood mononuclear cell (PBMC) derived HIV-1 JR-CSF viral supernatant was stored in separate aliquots of equal volume in order to compare the sensitivity of the Quantitect qRT-PCR viral load assay and the SG-PERT reverse transcriptase activity qPCR assay in parallel. (B) The Quantitect qRT-PCR viral load assay and the SG-PERT reverse transcriptase activity qPCR assay was run in parallel with viral RNA eluate and HIV-1 supernatant serially diluted until the limit of detection for each assay was reached. Data shown as the average cycle threshold (Cq) values determined from two technical replicates at each dilution. The limit of detection was defined as the Cq value at which the linear range of the assay ended. Absolute quantification of HIV-1 particles was determined from a viral RNA standard curve run in parallel with the Quantitect qRT-PCR viral load assay.

**Supplementary Figure 6:**
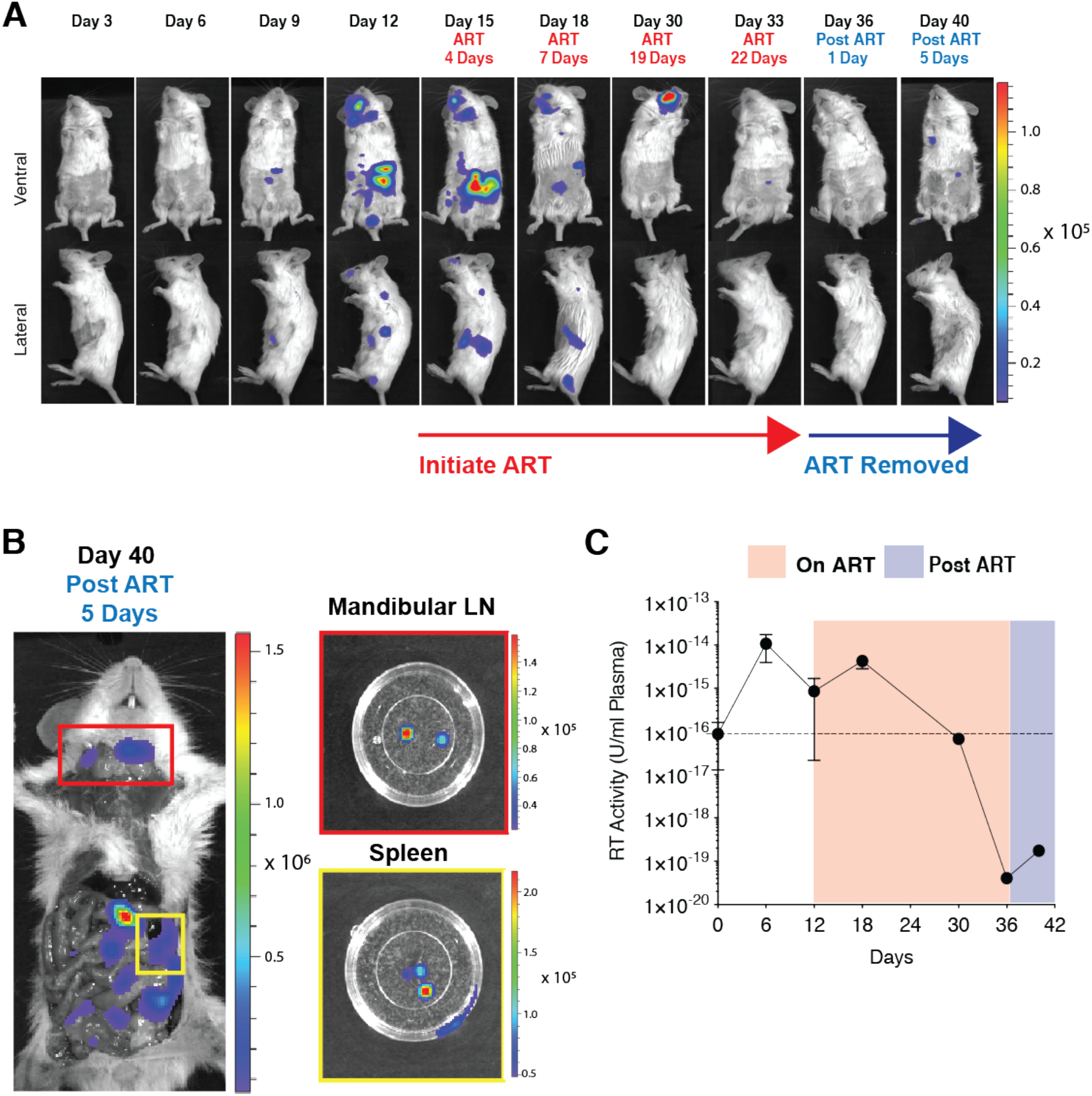
Longitudinal non-invasive bioluminescent imaging of HIV-1 acute infection, suppression, and recrudescent infection in the Hu-BLT mouse group placed on cART 12 days post-infection. (A) Bioluminescent imaging of spreading infection of Hu-BLT Mouse #3 infected with 1 x 10^6^ IUs of Q23.BG505.Nluc T/F reporter virus and placed on a daily cART regimen comprised of daily i.p. cART injections of Truvada and Isentress 12 days post-infection. (B) Whole animal ex vivo necroscopic analysis of rebounding infection in Hu-BLT Mouse #3 five days following cART cessation. (C) Plasma reverse transcriptase activity from Hu-BLT Mouse #3 over the course of the 40 day imaging period. Plasma reverse transcriptase activity in serum samples taken every six days over the course of the imaging period was measured via the SG-PERT reverse transcriptase activity assay and described as reverse transcriptase activity units / mL above endogenous uninfected background levels (dotted line).

